# Fluidized Bed Cathodes Enable High-Rate CO_2_ to Formate Production and In Situ Microbial Growth

**DOI:** 10.64898/2026.10.30.763118

**Authors:** Basem Zakaria, Antonio Lanzilotti, Eric Sundstrom, Sara Tejedor-Sanz

## Abstract

Electrochemical CO_2_ Reduction Reaction (eCO_2_RR) to formate offers a sustainable and promising route for storing renewable electricity in a microbial energy substrate. Conventional CO_2_ electrolyzers rely on flat electrodes with limited catalytic surface area, mass-transfer constraints, and electrolytes that are incompatible with microbial growth, preventing direct coupling of electrochemical formate production with biological conversion. Here, we developed a biocompatible CO_2_ electrolysis configuration that uses fluidized particles of Sn, Bi, and In as electrocatalysts to enhance CO□-to-formate conversion while supporting microbial utilization in the same chamber. The fluidized configuration enabled high formate production rates up to 0.24 mmol h□¹ cm□² with Faradaic efficiencies of 86–92%, representing a two-fold improvement over static electrodes. The system also achieved a carbon conversion efficiency of 42%, approaching the theoretical limit for co-generation of formate and bicarbonate. Importantly, electrolysis was performed in a biocompatible medium that supported direct in situ cultivation of *Cupriavidus necator* within the cathode chamber using only electricity and CO_2_/O_2_ as fed substrates. *C. necator* continuously consumed electro-generated formate for 5-6 days under aerobic conditions, demonstrating integrated CO_2_-to-biomass conversion. This work establishes a fluidized electrocatalytic platform for simultaneous electrosynthesis and microbial upgrading in a single reactor, providing a scalable strategy for renewable carbon fixation and sustainable biomanufacturing.

**TOC GRAPHIC:** 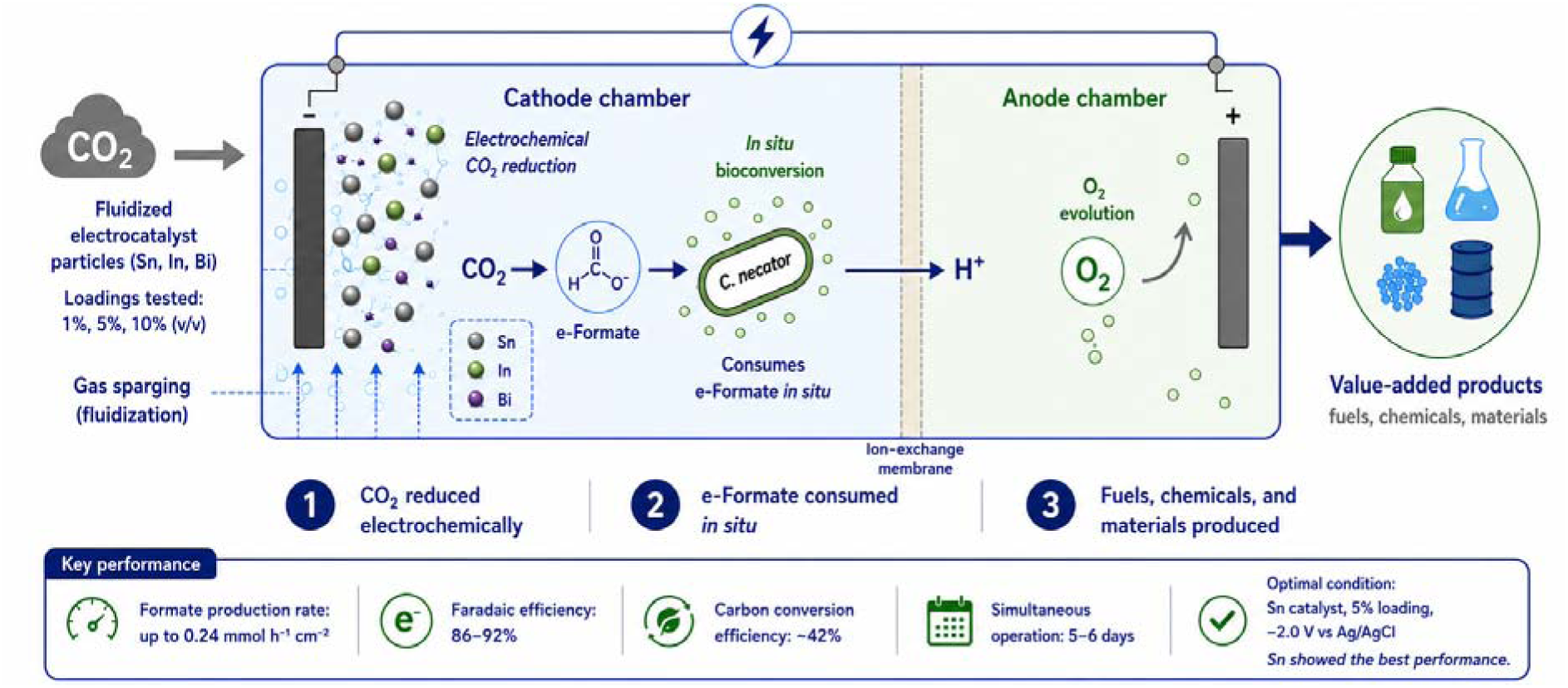

## INTRODUCTION

Converting CO_2_ into useful chemicals and fuels offers a route to close the carbon cycle while generating valuable commodities ^1–3^. Among such approaches, Electrochemical CO_2_ Reduction Reaction (eCO_2_RR) has gained significant attention as it enables CO_2_ reduction to organic molecules using renewable electricity, providing a carbon-neutral route to value-added products and generating reduced intermediates for further upgrading ^4–7^. Among the various eCO_2_RR pathways, reduction of CO_2_-to-formate has emerged as a particularly attractive route ^4–7^. Formate can be produced with high faradaic efficiency and selectivity, and it possesses high aqueous solubility and can serve as both a hydrogen storage carrier and a microbial feedstock ^8–10^. It can support the growth of diverse microorganisms and be further upgraded into higher-value multicarbon compounds, making it an appealing target product for CO□ electroreduction ^11^.

Recent advances in catalyst development have significantly improved the selectivity and efficiency of CO_2_-to-formate conversion. In particular, post-transition p-block metals such as tin (Sn), indium (In), and bismuth (Bi) exhibit exceptional catalytic activity and product selectivity towards formate while suppressing the competing hydrogen evolution reaction ^12–16^. These catalysts typically achieve formate faradaic efficiencies of 70–95% in aqueous bicarbonate-based electrolytes at cathodic potentials between −1.0 and −2.2 V vs. RHE (reference H_2_ electrode) ^17–21^. Beyond catalyst material, cathode and reactor design play a major role in system performance for eCO_2_RR. Flat cathodes provide relatively small surface areas and suffer from CO□ transport constraints, requiring large overpotentials to drive the reaction ^22–25^. In a static setup, there is also poor contact between the gaseous CO□, the liquid electrolyte, and the solid electrode, which further limits performance ^26–29^. Inclusion of microbes further exacerbates this dynamic, necessitating high surface area for efficient distribution of electrons and eCO_2_RR outputs to microbial cultures. For any scalable electrochemical reactor, especially in a bioelectrochemical context, maintaining a high electrode surface area relative to the reactor volume is critical ^26,30^. To overcome limitations of static electrodes, fluidized-bed electrodes have been proposed as alternative geometry to expand the catalytic surface and enhance mass transport in a biological context ^30,31^. In a fluidized-bed system, particulate catalysts are suspended and mixed (by gas bubbling, electrolyte recirculation or stirring), creating a dynamic 3D porous electrode through which reactants and ions can permeate freely. This strategy improves CO□ distribution, eliminates stagnant zones or clogging that can occur in packed beds, and increases catalyst– electrolyte contact ^26,32^.

Electrochemical CO□ reduction typically yields only short-chain C1–C3 products (e.g. CO, formate, methane) with mixed selectivity, while struggling to synthesize more complex multicarbon compounds ^33–35^. By contrast, autotrophic microbes can biosynthesize multicarbon with high yield and sensitivity from single carbon substrates ^9,36^. In particular, *Cupriavidus necator* can naturally consume formate as a sole carbon and energy source via formate dehydrogenases, entering central metabolic pathways or engineered assimilation routes for production of proteins, biopolymers, and other complex organics via aerobic biosynthesis ^37–39^. When compared to hydrogen as an alternative bioelectrochemical intermediate, delivery of formate as a soluble substrate unlocks substantial improvements in mass transfer and process safety, generating a highly promising platform for biomanufacturing from CO_2_ and renewable electrons ^40^.

For decoupled approaches separating CO_2_ electrolysis and bioconversion, electrolytically generated formate must be harvested, concentrated and transferred to a second bioreactor for microbial conversion ^41^. Formate accumulation may alter catholyte conductivity and induce anodic decomposition, reducing the process yield ^15,42^. Biochemical formate conversion results in significant generation of CO_2_ as a metabolic bioproduct, alongside accumulation of water in the bioreactor during fed-batch operation. Recycling of CO_2_ to the electrolyzer and removal of water to boost product titers results in energy losses, high infrastructure costs, and high overall system complex. Co-location of microbes in the same cathode compartment resolves these issues by enabling continuous in-situ reduction of generated CO_2_, eliminating water accumulation in the bioreactor, and utilizing biology to mitigate formate accumulation, greatly simplifying system design ^40,41^. This complementary relationship - rapid heterogeneous catalysis yielding simple molecules coupled to highly selective biological synthesis of complex products, has motivated the development of hybrid CO□ conversion platforms that integrate electrochemistry with biotechnology ^41,43^.

Many high-performance eCO□RR cathodes require strongly alkaline or purely inorganic electrolytes (e.g. concentrated KOH, carbonate buffers) that are lethal or inhibitory to bacteria ^36^. As a result, previous efforts to couple CO□ electrolysis with biological conversion have generally employed decoupled configurations ^40,41,44^. These challenges motivate the development of truly integrated systems in which electrochemical formate generation and microbial upgrading occur concurrently under compatible conditions.

In this work, we introduce a novel sandwich dual□chamber CO_2_ electrolysis platform designed for microbial assimilation of the in situ-produced formate. The core of the system is a fluidized-bed cathode of suspended catalyst particles (Sn, In, Bi) circulating in the catholyte, developed to be fully biocompatible to support *C.necator* growth. A key design feature of this platform is the deliberate use of a biocompatible catholyte and catalyst environment that enables electrochemical CO_2_ reduction to proceed under conditions suitable for microbial growth, rather than requiring separation between electrolysis and bioconversion. The fluidized-bed cathode further provides a dynamic, high-surface-area catalytic interface that improves CO_2_ mass transfer and catalyst-electrolyte contact while maintaining a physically accessible environment for in situ microbial formate assimilation. This approached constitutes a fully integrated biohybrid reactor in which formate is both produced and assimilated in one place, obviating any intermediate separation.

## MATERIALS AND METHODS

### Electrochemical cell configuration

Electrochemical experiments were conducted in a plexiglass sandwich-type H-cell reactor. The reactor consisted of two compartments (catholyte and anolyte), each with an effective working volume of approximately 30 mL, separated by an alkaline anion exchange membrane (Sustainion X37-50 Grade T, Dioxide Materials, USA) with a total surface area of 12.5□cm^2^. Before using the AEM membrane, pre-treatment was performed as described by the supplier by soaking in a bath of 1 M KOH for 24 hrs.

The cathode chamber housed a flat carbon fiber paper coated with a bismuth catalyst (Dioxide Materials, USA) that served as both current collector and flat catalytic surface. In addition, metallic microparticles of tin (Sn), bismuth (Bi), or indium (In) were suspended in the catholyte to form a fluidized electrocatalyst bed. Sn, Bi, and In particles were obtained commercially (−100 mesh, ThermoFisher Scientific, USA). Fluidization of the particles was maintained by continuous magnetic stirring combined with gas sparging. Particle loadings were adjusted to 1%, 5%, or 10% (v/v-net catholyte volume), depending on the experiment. An Ag/AgCl reference electrode (3M KCl, MF-2056, BASi, USA) was placed within 1 cm to the cathode current collector to minimize ohmic losses. The anode chamber consisted of a titanium electrode coated with a 10 µm layer of ruthenium-iridium (American Elements, USA) positioned parallel to the cathode. Both anode and cathode had an identical geometric area with 2 cm^2^. The cell was assembled using silicone gaskets to ensure gas-tight operation. All experiments were conducted at 30 °C, with the electrochemical reactors operating in an incubator. Continuous stirring was provided in both compartments using magnetic stir bars to maintain homogeneity and promote particle fluidization in the cathode chamber.

Electrochemical control and data acquisition were carried out using a multi-channel Biologic potentiostat (VMP-3e, BioLogic, USA) operated with EC-Lab software. Chronoamperometric experiments were performed by applying constant cathode potentials of –1.2 V, –1.5 V, or –2.0 V vs Ag/AgCl, depending on the experimental condition. Each experiment was operated continuously for up to 120 h, unless otherwise noted. Current density was calculated based on the projected geometric surface area of the flat cathode electrode (2 cm^2^). All potentials in this study are reported to Ag/AgCl in saturated KCl (0.197 V vs. standard hydrogen electrode (SHE)) unless stated otherwise.

### Electrolytes and media composition

Two different catholytes were used: (i) a conventional electrochemical electrolyte and (ii) a biocompatible growth medium. The conventional electrolyte consisted of 0.5 M KHCO_3_ prepared with ultrapure deionized water (18.2 MΩ·cm). The solution was purged with CO□ for at least 30 min prior to experiments to ensure saturation and pH stabilization. The biocompatible medium was formulated to support microbial growth while maintaining sufficient ionic conductivity for electrochemical operation. The composition of the biocompatible medium is provided in the Supplementary Information (**Table S1 and S2**). The initial pH of the catholyte was adjusted to 6.8 using HCl or KOH and monitored periodically throughout the experiments. The anolyte consisted of 0.5 M KHCO□ and was recirculated continuously using a peristaltic pump (L100-1S-2, Langer Instruments, USA) from a 300□mL reservoir at a flow rate of 1□mL min^-1^.

### Gas supply

High-purity CO□ gas (≥99.99% purity; Linde, USA) was continuously supplied to the cathode compartment through a gas sparger at a controlled flow rate of 15 mL min ¹ through a diffuser placed at the bottom of the cathode chamber (unless otherwise stated). Gas flow rates were regulated using a gas flow meter (Dakota Instruments, USA). The cathode outlet was connected to a gas flow meter and gas sampling bags for downstream gas analysis. The anode compartment was purged continuously with high purity nitrogen (99.99%) to flush O_2_ out of the anode chamber. All gas lines were equipped with sterile 0.2 µm filters when microbial experiments were performed.

### Biohybrid operating conditions

*Cupriavidus necator* H16 (ATCC 17699) was used as the model organism for biohybrid experiments. Acclimation of *C. necator* to the biocompatible medium and electrocatalyst particles was performed in shake flasks under aerobic conditions. Cultures were grown in biocompatible medium supplemented with glucose (5 g L□¹; **Table S1**) in the presence of electrocatalyst particles at different loadings (1%, 5%, and 10%). Flasks were incubated at 30 °C and agitated at 200 rpm. To determine the optimal aerobic gas composition for *C. necator* in the biohybrid system, growth experiments were first conducted in biocompatible medium supplemented with glucose (5 g L□¹) with varying CO□:O□ratios. This experiment was performed in 0.5 L automated gas fermentation reactors (bioXplorer 400P, HEL Ltd., UK) that allowed to continuously feed different gas compositions in parallel. The details of the bioreactor configuration are described in supplementary information **(Text S1).** Cell growth was monitored by measuring cell density to identify the gas composition that supported the highest growth rate. For the biohybrid experiments in the electrochemical cell, pre-cultures were grown aerobically in shake flasks using biocompatible medium supplemented with formate (30 mM) until mid-exponential phase. Cells were harvested by centrifugation, washed, and inoculated into the cathode chamber at an initial optical density of OD□□□ ≈ 0.1. In all experiments cell growth was monitored by measuring optical density at 600 nm (OD□□□) using a UV–Vis spectrophotometer (Genesys 150, ThermoFisher Scientific, USA).

### Analytical techniques

Liquid samples (500 µl) were periodically withdrawn from the cathode chamber using sterile syringes and filtered (0.2 μm) prior to analysis. Formate concentrations were quantified using HPLC (1260 infinity II, Agilent Technologies, USA) equipped with Bin pump (G7112B), diode array detector (G7115A), and refractive index detector (G7162A). 4 mM H_2_SO_4_ was used as a mobile phase. Calibration curves were prepared using sodium formate standards over a relevant concentration range. Gaseous products were collected in gas sampling bags connected to the cathode outlet and analyzed using gas chromatography (microGC 990, Agilent Technologies, USA) equipped with dual channel thermal conductivity detectors (TCDs). GC configured with two channels; a 20 m Molsieve 5Å column with argon as the carrier gas and 10 m PoraPLOT Q column with hydrogen as the carrier gas. The injection temperature was set at 80 °C, the MS5A was set to 100°C and the PPQ to 80 °C.

### Performance metrics and statistical analysis

Formate selectivity, rate of formate production, carbon conversion efficiency (CCE), and energy consumption per mole of formate (P*_formate_*) were calculated to evaluate the performance of CO_2_-to-formate conversion efficiency. Formate selectivity is reported as coulombic efficiency (CE), representing the fraction of electrons directed toward formate formation. More details of the metrics calculations are provided in the Supporting Information **(Text S2).**

## RESULTS AND DISCUSSION

### Effect of Cathode Applied Potential on CO_2_-to-Formate Conversion

We first aimed to evaluate the electrochemical conditions under which formate could be efficiently produced in our fluidized bed configuration. The cathode applied potential (E_cathode_) for CO_2_-to-formate conversion can significantly influence the reaction’s efficiency, selectivity, and competitive side reactions including the hydrogen evolution reaction (HER) ^45,46^. A wide range of cathode potentials have been previously reported for CO_2_-to-formate conversion utilizing flat cathode electrodes coated with different electrocatalysts^20,45,47^. Although various cathode electrode configurations, including static and flow cell setups, have been previously reported, this study will be the first to examine the behavior of fluidized electrocatalyst systems for eCO_2_RR under varying applied potentials.

The effect of applied E_cathode_ was investigated at three potentials ranges: −2 V, −1.5 V, and −1.2 V vs. Ag/AgCl. Three different systems were tested, each containing a different fluidized electrocatalyst composed of microparticles of Sn, Bi, and In, respectively, alongside a collector with high selectivity towards eCO_2_RR to formate. These applied potentials were chosen based on their previously demonstrated effectiveness for CO_2_ reduction reactions using these electrocatalysts ^20,45,48,49^. In our testing, eCO_2_RR to formate efficiency increased exponentially with decreasing cathode potential under all conditions examined. Using a static flat electrode at − 2 V (carbon electrode with deposited Bi electrocatalyst) resulted in a high formate production rate of 0.12 mmol hr^-1^ cm^-2^ with a maximum *CE* of 70% over 120 hours of operation, as compared to an E_cathode_ of −1.5 V (0.07 mmol hr^-1^ cm^-2^ and *CE* 59%) and −1.2 V (0.02 mmol hr^-1^ cm^-2^ and *CE* 38%) (**Fig. 1**). By contrast, a significant increase in performance was observed when adding fluidized electrocatalysts with the flat cathode electrode acting as both a current collector and electrocatalyst. Although the flat cathode electrode was composed of carbon paper coated with a Bi catalyst, known for efficient CO□-to-formate conversion, its performance was significantly lower than for the equivalent fluidized system, validating the impact of the fluidized particles in our configuration.

**Figure 1.**
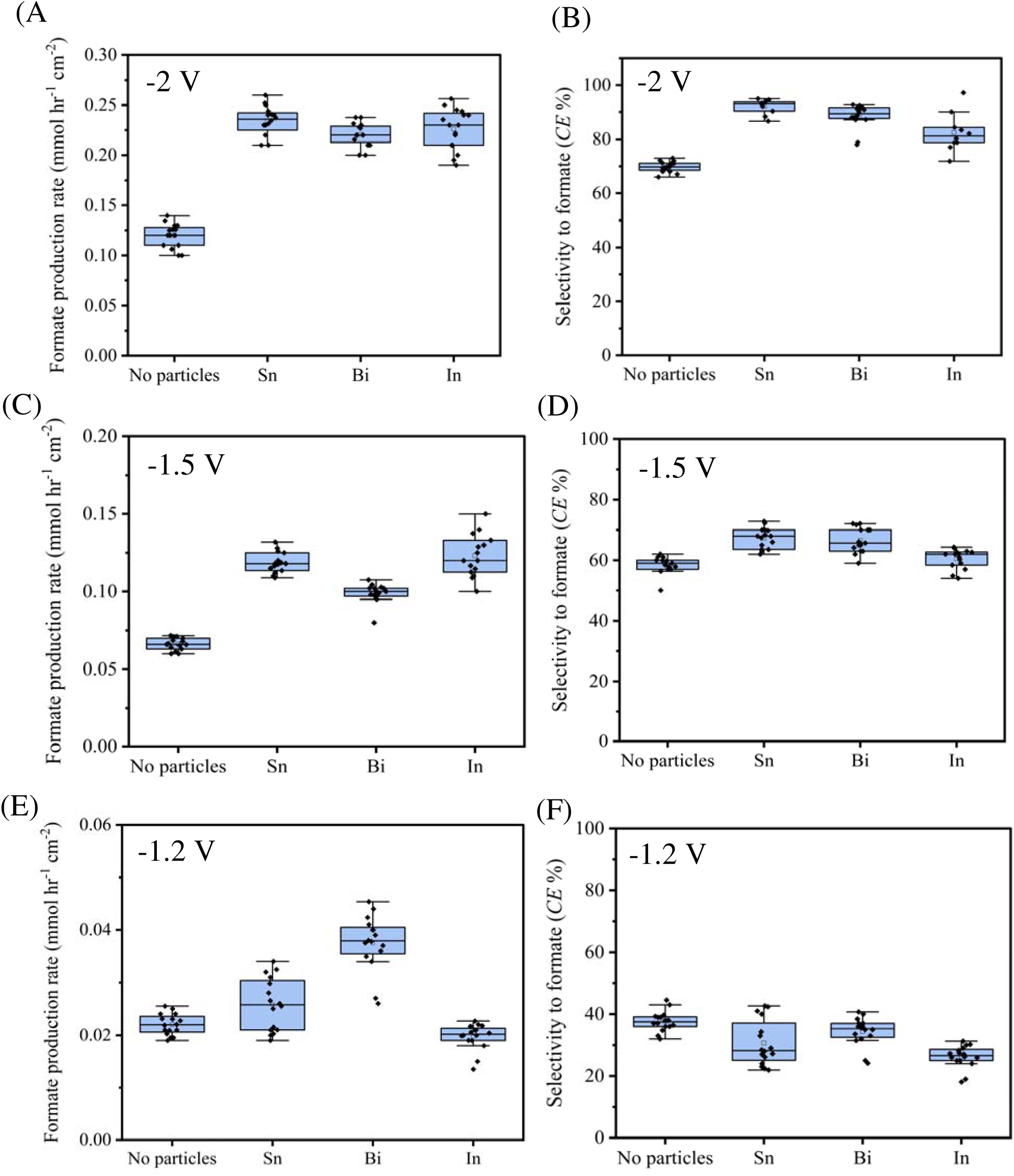
Effect of cathode applied potential on CO_₂_-to-Formate performance. (A-B) at −2 V, (C-D) at −1.5 V, and (E-F) at −1.2 V vs. Ag/AgCl showing corresponding formate production rates and *CE%*. All the catalysts were used at 5% (v/v-catholyte volume) loadings. Each condition was tested in duplicate, and each data point represents a sample taken at a steady state conditions.

Applying a potential of −2 V achieved the highest formate production rate of 0.22-0.24 with a *CE* of 86-92%. When the applied potential was adjusted to −1.5 V, the formate production rate decreased to an average 0.09-0.12 mmol hr□¹ cm□² with *CE* 62-68%, and at −1.2 V, the rate further declined to 0.02-0.04 mmol hr□¹ cm□² and *CE* 26-35% (**Fig. 1**). In agreement with the literature, our fluidized electrocatalyst configuration demonstrated that negative potential enhances both the CO_2_ reduction reaction rate and *CE* to formate ^20,45,48,49^. Current density consumption was also observed to increase with more negative applied potentials across all experimental conditions (**Fig. 2A**). Without the addition of fluidized electrocatalyst particles, current density significantly increased from 3.1 mA c□ ² at −1.2 V to 6.0 mA cm□² at −1.5 V, reaching 9.5 mA cm□² at −2 V. The addition of fluidized electrocatalyst particles (Sn, In, and Bi) further enhanced the current density at all tested potentials. Current densities increased to 4.0-5.9 mA cm□^2^ at −1.2 V and 8.2-10.7 mA cm□^2^ at −1.5 V, with the most substantial enhancement observed at −2.0 V, where current densities reached 14.5-15.6 mA cm□^2^. These results show that the formate production rate and selectivity towards formate were significantly enhanced by the addition of any of the fluidized electrocatalysts tested.

**Figure 2.**
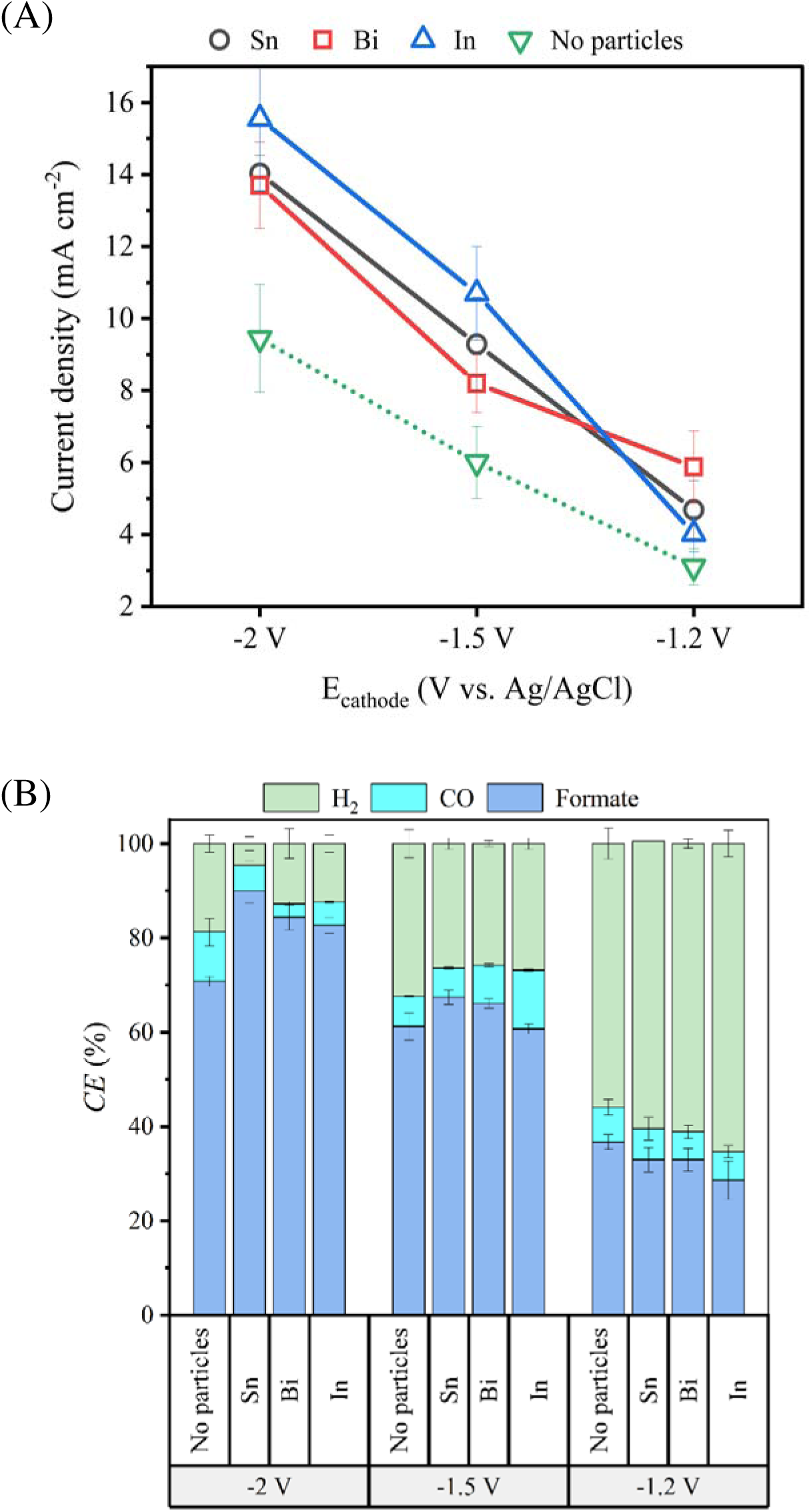
Effect of cathode potential on current density and *CE* for the three electrocatalysts tested. (A) Current density, and (B) *CE* under different applied potentials and electrocatalysts. All the catalysts were used at 5% (v/v-catholyte volume) loadings. Experiments were performed in duplicate, and results are reported as mean ± standard deviation.

More negative applied potential not only increased the rate of formate production but additionally enhanced selectivity towards the formate (**Fig. 2B**). Specifically, at −2 V, formate selectivity was maximized with *CE* 70-92%, while HER was minimized in the range of *CE* 5-19%, indicating a shift in electron utilization favoring CO_2_ reduction towards formate. A notable decrease in formate selectivity occurred as the potential was switched to less negative values (−1.5 V) with *CE* of 60-68%, and a shift in the selectivity towards HER with *CE* of 26-32%. By further applying lower negative potentials (−1.2 V), HER became the dominant electron sink with CE of 56-65%, substantially limiting formate production within CE of 30-38%. It is noteworthy that the overpotentials for HER and CO□ reduction to formate are very similar ^18,20^. Although thermodynamics indicate that both reactions can occur, the selectivity is largely driven by the kinetics of intermediate formation and catalyst specificity, making mechanistic and kinetic factors the key influences in CO□ reduction to formate over HER. Overall, the trend observed across all conditions suggests that applied potential is a critical factor influencing the efficiency of CO_2_-to-formate conversion. This higher potential enhances availability of electrons, facilitating the effective reduction of CO_2_ to formate. Moreover, energy consumption analysis showed that fluidized catalyst configurations achieved lower energy input per mole of formate compared to flat electrodes, particularly at −2 V applied potential. A detailed energy efficiency analysis is provided in the Supporting Information **(Text S3, Fig. S1)**. Multivariate principal component analysis (PCA) further confirmed strong correlations between applied potential, current density, and formate production rate, highlighting the dominant influence of operating potential on system performance (**Text S4, Fig. S2**). Given that −2 V maximized both the formate production rate and selectivity, this potential was chosen for subsequent experiments.

### Role of the fluidized electrocatalyst material in CO_2_-to-formate conversion

Next, we investigated the impact of three materials as fluidized electrocatalysts on CO□-to-formate conversion efficiency: Sn, Bi, and In at different applied potentials. Among the electrocatalysts tested, Sn particles exhibited the highest formate production rate, achieving 0.24 mmol hr^-1^ cm^-2^ and a high CE of 92% (**Figure 3**). Although performance of Bi and In particles did not reach that of Sn, they also showed high formate production rates and high selectivity to formate, achieving 0.22 mmol hr^-1^ cm^-2^ and *CE* 86%, and 0.23 mmol hr^-1^ cm^-2^ and *CE* 82% respectively when operated for 120 hrs. In the flat cathode electrode system without fluidized electrocatalysts, the formate production rate was significantly lower at 0.12 mmol hr^-1^ cm^-2^ and *CE* of 70%. Results from the other tested cathode applied potentials (−1.5 V and −1.2 V) are shown in the Supplementary Information **(Fig. S3)**. Current density also increased in the presence of the fluidized electrocatalysts as compared to the flat electrode alone. The highest current density of 16 mA cm^-2^ was achieved with In electrocatalysts, followed by 14 mA cm^-2^ with Sn and Bi particles, with the flat electrode system reaching only 9.5 mA cm^-2^ (**Fig. 2B**). The fluidized electrocatalysts not only improved mass transfer of CO_2_ to the catalysts but also expanded catalytic capacity for CO_2_-to-formate conversion across the cathode working volume. We hypothesize that slight differences in the performance among electrocatalysts may be due to their distinct electronic configurations and surface interactions with CO_2_ during the CO_2_-to-formate process ^50^. Given the high cost and limited availability of In and Bi, the Sn particles stand out as a promising candidate for CO_2_ reduction to formate due to both their affordability, and their efficient electrochemical performance as shown in this study ^51,52^.

**Figure 3.**
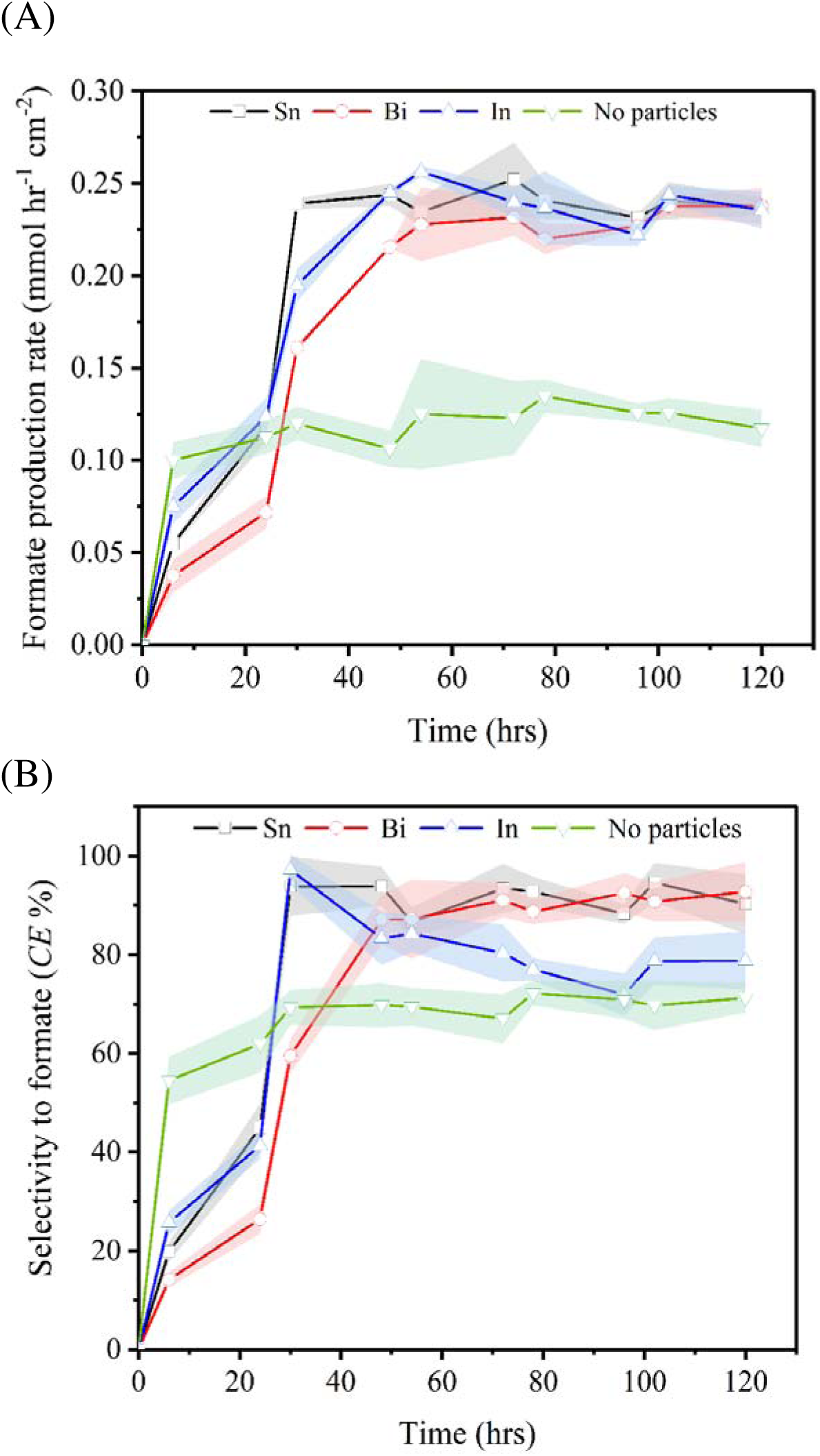
Influence of electrocatalyst material on CO_2_-to-Formate at −2 V vs. Ag/AgCl. Results from the other tested cathode applied potentials (−1.5 V and −1.2 V) are shown in the Supplementary Information. All the catalysts were used at 5% (v/v-catholyte volume) loadings. Experiments were performed in duplicate, and results are reported as mean ± standard deviation (shown with a shaded area).

Formate production in the fluidized system gradually increased over the first 24 hours before reaching a plateau, where it remained stable for the following 96 hours. This behavior differs from the flat electrode where the formate production rate remained stable throughout the operation. This gradual increase in formate production is attributed to the stabilization of fluidized particle dynamics, electrochemical polarization, and gas–liquid mass transfer. In fluidized systems, the effective electroactive surface area evolves over time as particle–collector interactions, local current distribution, and CO_2_ availability reach steady-state conditions ^53^.

### Effect of electrocatalyst loadings on eCO_2_RR in the fluidized system

As aforementioned, the fluidized catalyst increased the effective surface area available for eCO_2_RR, where the interaction between the particles and current collector is critical to enhance the catalytic capacity of cathode chamber. By optimizing the load of the fluidized catalyst, we hypothesize the number of available active sites increases, allowing for more CO□ molecules to interact with catalytic surfaces and the collisions particles-current collector enhance. To evaluate the impact of electrocatalyst loading on the performance of the CO_2_ reduction system, experiments were conducted using different loadings of Sn fluidized electrocatalysts in the cathode chamber at 1%, 5%, and 10% (v/v-catholyte volume) **(Fig. S4)**. These conditions were compared to a baseline without Sn fluidized particles (0%) to determine whether increasing the particulate surface area enhances the catalytic performance. Without addition of particles (0%), the formate production rate was 0.12 mmol h□¹ cm□², with a CE of 70% and a current density of 9.5 mA cm□². The addition of 1% catalyst loading improved the formate production rate to 0.16 mmol h□¹ cm□², the CE to 83%, and the current density to 10.5 mA cm□², demonstrating better performance compared to the baseline. Although the 1% catalyst loading outperformed the 0% condition, it exhibited the lowest efficiency among the tested catalyst loadings, as higher particle concentrations (5% and 10%) provided further improvements in catalytic activity and *CE*. Thus, a low catalyst loading was insufficient to exploit the distinct catalytic properties of the catalyst. This may be due to the limited active surface area and reduced probability of particle collision and electron transfer with the current collector and the CO_2_ molecules, which constrained the catalytic reactions.

Upon increasing the catalyst loading to 5%, a substantial improvement in performance was observed, and the highest rate of formate production and selectivity towards formate was achieved. Formate production rate increased significantly to 0.24 mmol hr^-1^ cm^-2^, achieving a *CE* of 91% with Sn electrocatalyst. Furthermore, the current density was enhanced 14 mA m^-2^ with a 5% of catalyst loading. This increase in catalytic performance can be attributed to the enhanced availability of active sites, as higher particle density in the fluidized state promotes more frequent and efficient contact between catalyst surface, CO_2_ molecules and current collector, boosting reduction reaction efficiency. Additionally, the increased catalyst concentration facilitates electron transfer processes within the fluidized catalysts. By further increasing the catalyst loading to 10%, there was no performance enhancement over the 5% loading. This plateau in performance suggests that at higher loadings, factors such as particle agglomeration and mass transport limitations begin to offset the potential benefits of additional active sites. Also, at high catalyst concentrations, inter-particle interactions could hinder optimal dispersion, potentially leading to uneven current distribution and limited access of CO_2_ to the active sites. Thus, the 5% loading of Sn particles appears to be optimal concentration for achieving high formate selectivity and production efficiency with our configuration.

### Carbon conversion efficiency of the system

While *CE* assesses how efficiently the supplied electrons are utilized in the target electrochemical reaction for CO_2_ reduction to formate, carbon conversion efficiency (*CCE*) emphasizes the effectiveness of converting the CO_2_ as a C-source into the desired carbon-based product – in this case formate. The *CCE* was calculated as the ratio of CO_2_ converted to formate relative to the total CO□ supplied to the system (**Fig. 4**). In order to evaluate the *CCE* of the system, a carbon balance of the entire process was performed using the Sn catalyst due to its superior overall performance.

**Figure 4.**
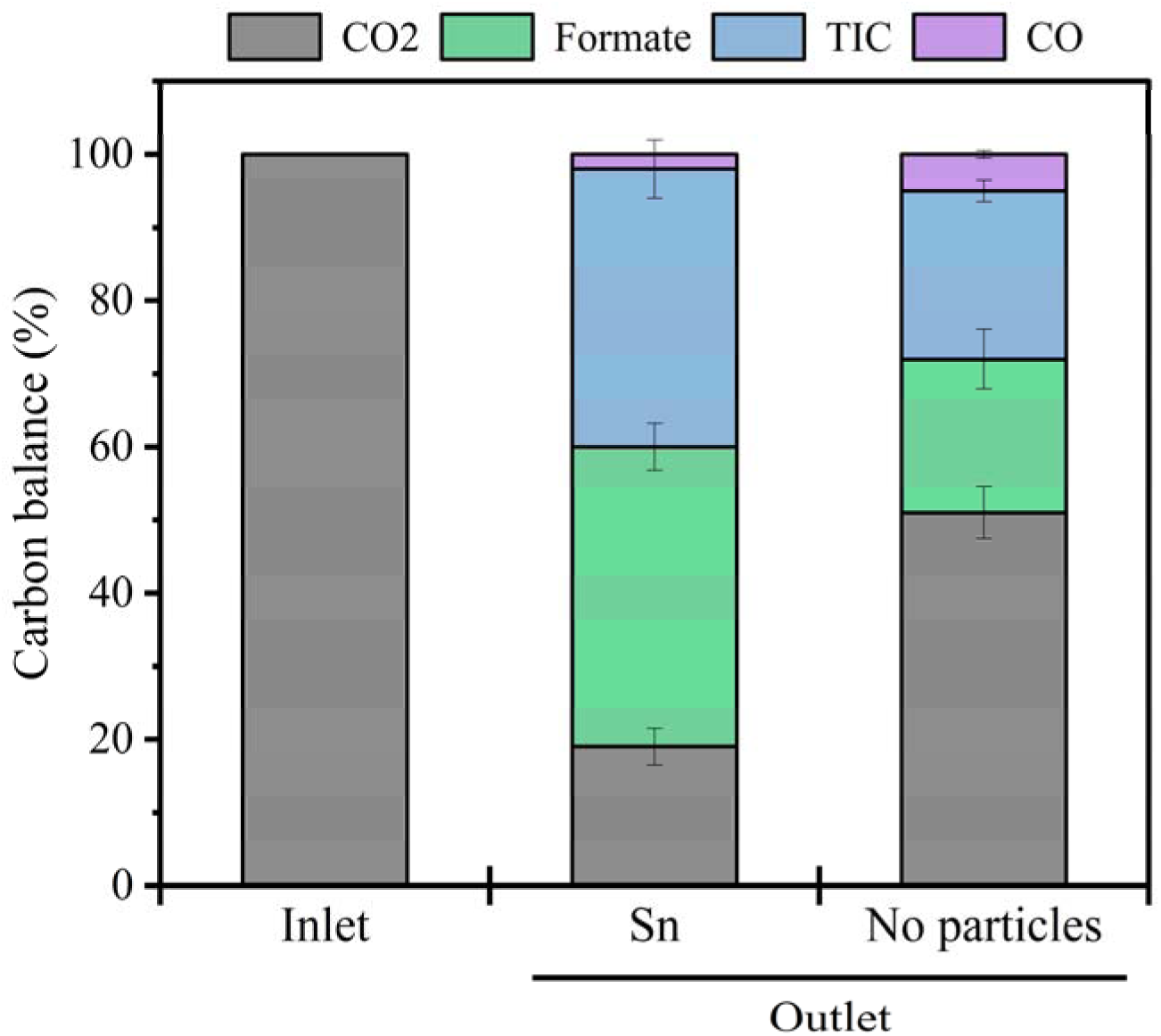
Carbon Balance and Carbon Conversion Efficiency (CCE) of the eCO2RR in the fluidized system. Sn catalyst with 5% (v/v-catholyte volume) loading at −2 V vs. Ag/AgCl was used. Experiments were performed in duplicate, and results are reported as mean ± standard deviation.

A low *CCE* was observed when using the flat electrode without Sn catalyst, with only 21% of supplied CO_2_ converted to formate. Of the total CO_2_ supplied, 51% remained unreacted and was released as off-gas, 23% was chemically flat as inorganic carbon in the form of bicarbonate (HCO□□), and 5% was converted to carbon monoxide (CO) (**Fig. 4**). The system demonstrated a significant improvement when Sn particles were added as fluidized catalyst, achieving *CCE* values exceeding 42% for formate production. Despite this increase, 18% of the CO_2_ remained unconverted and was released as off-gas, 38% was chemically flat as bicarbonate and 2% was converted to CO. This performance approaches the theoretical maximum *CCE* for CO_2_ to formate conversion, which cannot exceed 50% as the process requires 2 moles of CO_2_ to simultaneously produce formate and HCO ^-54^. A recent study reported a *CCE* of 41% for CO − to-formate conversion in a flow cell using pure KHCO_3_ as the catholyte. Notably, our study achieved a similar *CCE* of 42% marking the first instance of such efficiency being attained in a biocompatible medium within a sandwich H-cell configuration, and with fluidized particles. Additionally, the high bicarbonate produced on the cathode chamber in our system may have acted as a carrier to transfer hydroxide ions from cathode to anode stabilizing the catholyte pH ^55,56^. We attribute the high *CCE* achieved with the Sn catalyst to a highly specific catalytic activity and effective CO_2_ utilization on the reduction process, minimizing competitive side reactions including HER and CO production. The fluidized Sn catalyst may have also contributed significantly to high CO_2_ conversion rates by enhancing mass transfer and improving reaction kinetics.

An energy efficiency analysis of the CO_₂_-to-formate conversion process was performed by determining the energy consumption in kilowatt-hours per mole of formate under varying cathodic potentials and fluidized catalysts materials (see Supporting Information). The Sn and Bi fluidized system at −2 V, or the In at −1.5V, provided the best combination of low energy consumption and high formate selectivity, making them the best candidates for upscaling. Overall, the results demonstrate the critical impact of both potential and catalyst type on the energy efficiency of the system. Furthermore, to assess relationships between operating parameters and system performance, a correlation analysis was conducted using Spearman’s rank correlation coefficient (ρ), and principal component analysis (PCA) methodology (see Supporting Information).

### Performance of the system under aerobic and anaerobic environments

Results under abiotic conditions demonstrate that the biohybrid electrosynthesis system can efficiently and effectively convert CO□ into formate, a potential substrate to be consumed by various microbial species within the cathode chamber. Although the fluidized configuration demonstrated high selectivity and coulombic efficiency for CO□-to-formate conversion, the system was supplied with a pure CO□ fed in the absence of O□. Under these anaerobic conditions, the reactor environment might favor the growth of only anaerobic microbial species, which can potentially utilize the produced formate. However, many biotechnological platform organisms that can utilize formate to efficiently produce complex molecules including fats, proteins, polymers, and drop-in aviation fuels require O_2_ - including *Cupriavidus necator* and genetically modified species of *Escherichia coli*, *Saccharomyces cerevisiae* or *Pseudomonas putida* ^57^. To date, minimal published data exists examining the influence of an aerobic environment on eCO_2_RR system performance using a full biocompatible medium ^21,58^. Introducing O□ into the system could negatively impact the selectivity and coulombic efficiency for CO_2_-to-formate conversion, as oxygen reduction can be a significant competing reaction at the cathode (O_2_ + 4H□+ 4e□ → 2H_2_O), consuming electrons that would otherwise drive the reduction of CO_2_ to formate. In fact, the oxygen reduction reaction occurs at a less negative – and therefore more favorable - potential than the eCO_2_RR.

To assess the impact of O□ on CO□-to-formate conversion efficiency, we purged the system with a mixed gas composition of 25% air (5% O_2_) and 75% CO_2_ (**Fig. 5**). This gas composition was selected based on experiments conducted in bioreactors with varying O□:CO□ ratios, which demonstrated that this mixed gas composition achieves the highest growth rate of *Cupriavidus necator* **(Fig. S5).** Interestingly, O_2_ purging had only a minor impact on the rate and *CE* compared to the anaerobic environment. These trends were observed with both the fluidized Sn electrocatalyst and the flat electrode. In the fluidized system, the aerobic environment decreased the production rate and *CE* from 0.24 to 0.22 and 92% to 86% as compared to the anaerobic environment (**Fig. 5**). Similarly, the flat electrode experienced a mild decline of both rate and *CE* in the aerobic environment, where it also dropped from 0.12 to 0.11 and 70% to 68%. This decrease may result from the impact of O_2_ on CO_2_-to-formate efficiency leading to a slight decline in rate and selectivity. Our observations are in agreement with Hegher *et al* who observed that aerobic conditions of 3% had a minor impact on eCO_2_RR when using a 0.05 M NaHCO_3_ electrolyte solution ^21^. Overall, these results demonstrate that our CO_2_-to-formate electrosynthesis system could be biocompatible with both anaerobic and aerobic metabolism, highlighting the feasibility of coupling CO_2_-to-formate electrosynthesis with microbial utilization in diverse applications.

**Figure 5.**
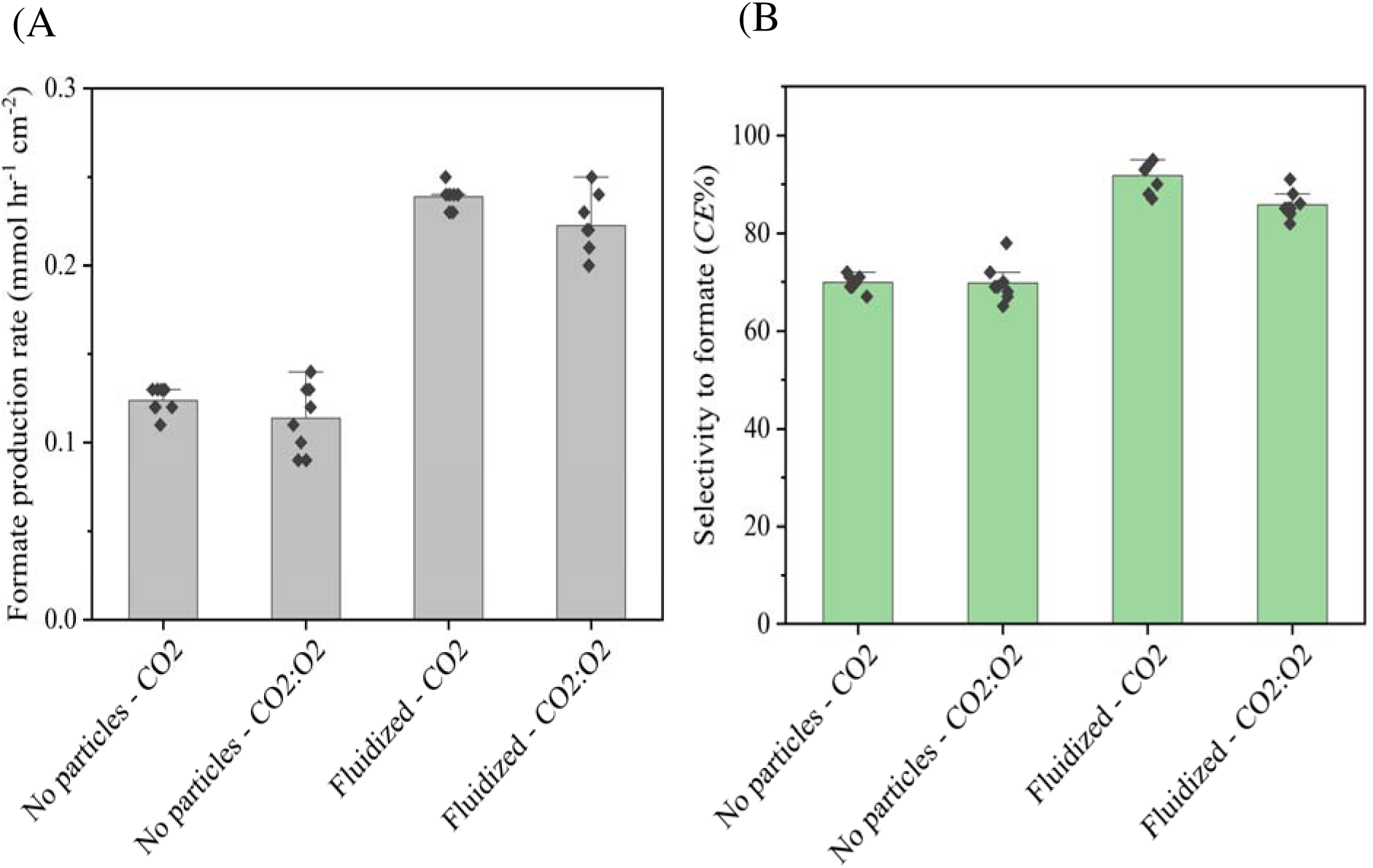
Influence of aerobic vs. anaerobic conditions on CO_₂_-to-Formate efficiency. For aerobic condition, 25% air (5% O_2_) and 75% CO_2_ gas mixture was used, and for anaerobic condition, CO_2_ was the only gas supplied. Sn catalyst with 5% (v/v-catholyte volume) loading at −2 V vs. Ag/AgCl was used. Experiments were performed in duplicate, and results are reported as mean ± standard deviation of the data points collected at steady-state.

### Performance with a biocompatible media versus a conventional electrochemical electrolyte

In CO_2_ electrolysis, potassium bicarbonate (KHCO_3_) or sodium bicarbonate (NaHCO_3_) at high concentrations are widely used as catholytes due to their effective buffering capacity and favorable ionic conductivity ^59–64^. However, due to high salinity and the presence of toxic constituents these media are generally non-compatible with microbial growth, presenting a barrier to coupling CO_2_ electrolysis with microbial utilization of formate in the same reactor. To facilitate this integration, it is essential to optimize the catholyte to support both efficient formate production and microbial viability. Thus, we compared the performance of a 0.5 M KHCO_3_ electrolyte and a biocompatible medium formulated to support microbial growth using the Sn fluidized electrocatalyst (**Fig. 6**). Details of the biocompatible medium composition are provided in the supplementary information **(Table S1)**. We observed that the KHCO_3_ electrolyte yielded a higher formate production rate of 0.3 mmol hr□¹ cm□², as compared to 0.24 mmol hr□¹ cm□² using the biocompatible medium. Despite the higher rate observed with KHCO_3_, there was no significant difference in *CE* between the two media with 93% in KHCO_3_ and 91% in biocompatible media, indicating that both supported the CO_2_-to-formate conversion with similar selectivity. The high CO_2_-to-formate efficiency observed in 0.5 M KHCO_3_ is attributed to its higher conductivity of 27.3 mS cm^-1^, compared to 7.4 mS cm^-1^ of the biocompatible media. This difference in the conductivity is due to the increased bicarbonate (HCO ^-^) concentrations, as higher HCO ^-^ levels enhance ionic conductivity. Additionally, the higher concentration of HCO ^-^ ions facilitate OH^-^transfer from the cathode to the anode through the anion exchange membrane, helping to maintain electroneutrality and stabilize the pH of both the anolyte and catholyte ^55,56^.

**Figure 6.**
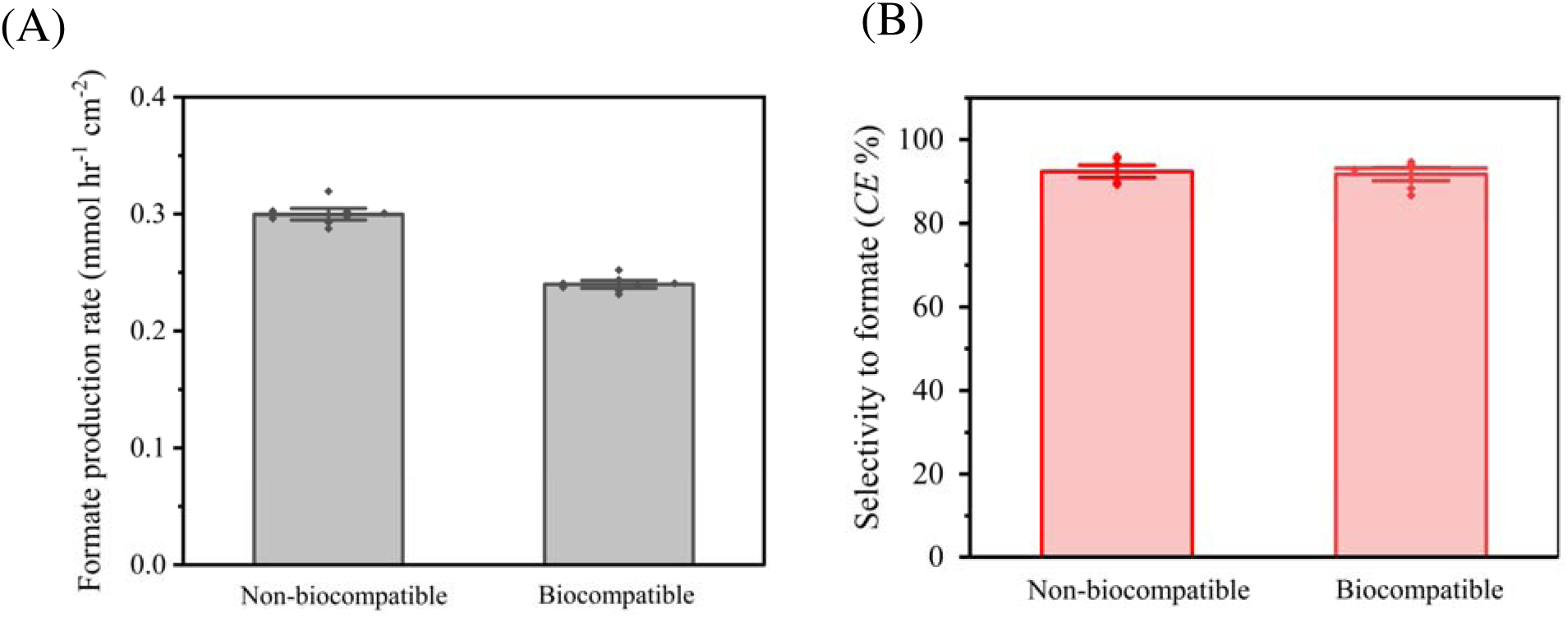
Comparison of biocompatible medium vs. KHCO_3_ electrolyte on CO_2_-to-Formate efficiency. Sn catalyst with 5% (v/v-catholyte volume) loading at −2 V vs. Ag/AgCl was used. Experiments were performed in duplicate, and results are reported as mean ± standard deviation of the data points collected at steady-state.

These findings suggest that while KHCO_3_ increases formate production rate, the biocompatible medium provides a viable alternative for systems requiring microbial growth, with a relatively small decline in performance trading off against the substantial benefits of the integrated configuration. Further optimization of biocompatible media could help narrow the gap in production rates while maintaining selectivity, creating a pathway for more efficient and sustainable biohybrid CO_2_ electrolysis systems.

### Biocompatibility of the Catholyte and the Fluidized Electrocatalyst Particles with Microbial Growth

After testing the formate selectivity and production rates using a biocompatible catholyte, we aimed to acclimate *C. necator* to grow in the biocompatible media and with the electrocatalyst particles. *C. necator* has long served as a benchmark organism for formate-supported and CO□-derived bioproduction due to its robust central metabolism, high growth rates on C□ substrates, and extensive characterization in both autotrophic and formatotrophic conditions ^65,66^. This allowed us to evaluate whether the biohybrid system could sustain continuous microbial growth under formate electrosynthesis conditions. To analyze the impact of the electrocatalyst on microbial growth and further optimize growth and eCO_2_RR under hybrid bioelectrochemical operation, we again evaluated variable particle loadings of 1%, 5% and 10%. The OD_600_ growth curves over three sequential batch transfers revealed clear evidence of acclimation of *C. necator* to the biocompatible media and electrocatalysts (**Fig. S6).**

Initially, the presence of Sn, Bi, or In particles imposed a growth inhibition manifested as lower OD and longer lag in the first transfer compared to a catalyst-free culture. By the 3^rd^ transfer, *C. necator* exhibited substantially improved growth rates, a shorter lag phase, and final high cell densities on most catalyst conditions (OD_600_ 2-2.5) indicating that the bacteria acclimated to the particle-induced stress over time. Among the materials tested, Sn showed to have the minimum impact on growth among all particle loads tested, which could be due to a lower propensity to leach toxic ions or less cell damage.

At 5% (w/v) particle loading, cultures with Sn, In, or Bi catalysts showed OD600 profiles comparable to or even exceeding the particle-free control, indicating high baseline biocompatibility of all three catalysts. Notably, the 5% Sn condition supported the highest cell density, with growth surpassing that of the particle-free control **(Fig S6)**. Increasing the catalyst loading to 10% (w/v) substantially suppressed growth with all three metals. This inhibition at higher loadings is likely due to increased cellular stress from the excess solids, greater metal ion toxicity through leaching of Sn, In, or Bi ions, or physical hindrances such as particle aggregation and crowding that can impede mass transfer and even cause cell damage. Overall, these results demonstrate the viability of culturing *C. necator* to high cell densities in a fluidized catalyst environment with a 5% particle loading, while excessive particle concentrations (10%) can negatively impact biomass production, likely through combined toxic and physical stress mechanisms.

### Biohybrid Integration of CO□ Electrolysis with In Situ Microbial Growth

To evaluate the integration of the in-situ bioconversion of formate with the electrosynthesis, *C. necator* was cultivated in the cathode chamber while performing CO_2_-to-formate electroreduction in the same compartment. Unlike previous systems that physically separated the electrosynthesis and bioconversion steps, here both processes are truly integrated in one chamber and simultaneous.

Before directly culturing *C. necator*, careful control of the formate production rate is needed to avoid formate toxicity due to the accumulation of this substrate. Thus, we deliberately operated the cathode at a mild potential (–1.2□V vs Ag/AgCl), inherently limiting the formate generation rate. Also, based on the acclimation results, Sn catalyst with 5% loading was used in the catholyte to enhance cell growth (**Fig. S6**). An abiotic control was first operated in the biohybrid fluidized configuration to establish formate production rates, which were maintained steadily at 0.05 mmol hr□¹ cm□² (42 mM) over time, confirming continuous CO□-to-formate conversion with lower formate toxicity levels (**Fig. 7A**). Previous studies demonstrated that formate concentrations on the order of 50–100 mM inhibit growth of *C. necator* and other formatotrophs ^66–68^. While using a lower potential sacrifices conversion rate, it maintains formate output in pace with cellular uptake, preventing accumulation. In our system, the steady-state formate levels in the abiotic reactor were below these inhibitory thresholds (**Fig. 7A**). By matching the production and consumption rates, the reactor effectively buffered the formate concentration and prevented any toxic buildup. To further enhance biocompatibility, an optimized loading of 5% Sn catalyst with 5% loading was used in the catholyte to enhance cell growth. Under these conditions, *C. necator* growth (OD600) reached 0.80 after 60 hrs with steady-state growth observed until 144 hrs of operation (**Fig. 7B**). Formate electrosynthesis and microbial growth was sustained simultaneously for ∼5–6 days, however growth from day three onward was slow or even stalled, potentially due nutrient depletion, waste accumulation, or longer-term release of metal ions. This long-term stability is noteworthy, as many bioelectrochemical systems struggle to maintain activity over multi-day timescales ^69,70^. Single chamber biochemical-integration demonstrates a clear advantage in this use case: the formate is utilized at the site of production, enabling a seamless flow of carbon from CO_2_ to formate and immediately into biomass. Biological consumption effectively maintained formate concentrations below the toxicity threshold for bioconversion (**Fig. 7B**), while simultaneously mitigating feedback inhibition of the electrochemical process. Further optimization of catholyte electrochemical conditions could prolong concurrent CO_2_ electroreduction and in-situ microbial cultivation and generate higher cell densities and bioproduct titers in a single reactor.

**Figure 7.**
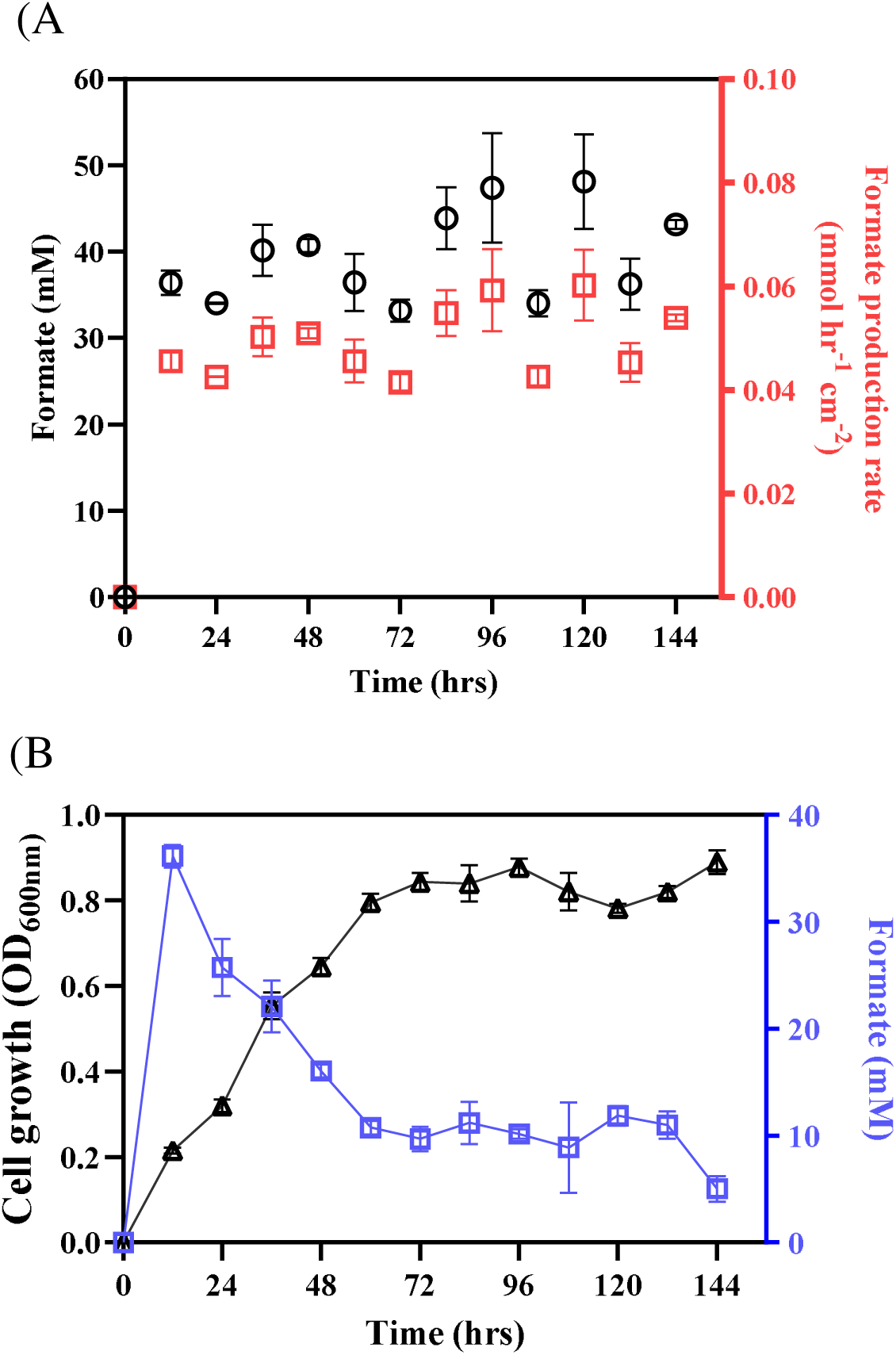
In-situ eCO_2_RR to formate and *C. necator* growth using fluidized Sn particles as cathode. (A) Abiotic condition test, and (B) Biohybrid integration of *C. necator* in the catholyte. Sn catalyst with 5% loading was used in the catholyte, and applied potential was set at −1.2 V vs. Ag/AgCl. Experiments were performed in duplicate, and results are reported as mean ± standard deviation.

## CONCLUSIONS

In this work, we developed a biocompatible fluidized-bed CO□ electrolysis platform that enables efficient formate electrosynthesis while supporting microbial growth within the same reactor. By systematically evaluating operating parameters, including applied potential, catalyst material (Sn, Bi, and In), catalyst loading, electrolyte composition, and gas atmosphere, we identified key design principles for maximizing □-to-formate performance while maintaining biological compatibility. The fluidized electrocatalyst configuration significantly outperformed conventional flat cathodes, achieving formate production rates up to 0.24 mmol h□¹ cm□² with Faradaic efficiencies of 86–92%, approximately twice that obtained with static electrodes. Optimal operation occurred at −2.0 V vs Ag/AgCl with ∼5% (v/v) catalyst loading, conditions that maximized catalytic surface area and mass transfer. Under these conditions, the system achieved a carbon conversion efficiency of ∼42%, approaching the theoretical limit for CO_2_-to-formate conversion in bicarbonate-based electrolytes. The reactor also maintained high selectivity and stable current in both anaerobic and oxygen-containing environments.

Importantly, the electrolysis was conducted in a biologically compatible medium that supported direct cultivation of *Cupriavidus necator* within the cathode chamber. The microorganisms continuously consumed electro-generated formate over several days of operation, demonstrating simultaneous electrosynthesis and microbial assimilation in a single reactor. These results establish fluidized-bed electrocatalysis as a promising platform for integrated electro-biological CO_2_ conversion and provide a foundation for scalable CO_2_-to-biomass and sustainable biomanufacturing systems.

## ASSOCIATED CONTENT

### Supporting Information

Biocompatible electrochemical media (catholyte) for *C. necator* growth (Table S1); trace minerals solution for biocompatible media (Table S2); bioreactor configuration (Text S1); performance metrics and statistical analysis (Text S2); energy efficiency and power input (Text S3); principal component analysis (Text 4); effect of electrocatalyst loadings on CO_2_RR in the fluidized system (Text S5); energy efficiency (Figure S1); multivariant analysis (Figure S2); influence of electrocatalyst materials (Figure S3); effect of catalyst loading (Figure S4); *C. necator* growth under aerobic conditions with different CO_2_:O_2_ mixtures (Figure S5); acclimation of *C. necator* growth in the biocompatible media with electrocatalysts (Figure S6).

## ACKNOWLEDGEMENTS

This work was supported by the U.S. Department of Energy, Office of Science, under the contract DEAC02-05CH11231 between DOE and Lawrence Berkeley National Laboratory, through a Laboratory Directed Research & Development (LDRD) Program within the Biological Systems and Engineering (BSE) Division and the Advanced Biofuels and Bioproducts Process Development Unit (ABPDU).

## CREDIT AUTHORSHIP CONTRIBUTION STATEMENT

**Basem Zakaria:** Writing – review & editing, Writing – original draft, Experimental work, Visualization, Validation, Formal analysis, Conceptualization. **Antonio Lanzilotti**: Visualization, Validation, Formal analysis**. Eric Sundstrom:** Writing –review & editing, Visualization, Validation, Conceptualization. **Sara Tejedor-Sanz:** Writing – review & editing, Writing – original draft, Resources, Project administration, Funding acquisition, Conceptualization.

## DATA AVAILABILITY STATEMENT

The data that support the findings of this study are available from the corresponding author upon reasonable request.

