## Supplementary Information for "Fluidized Bed Cathodes Enable High-Rate CO_2_ to Formate Production and In Situ Microbial Growth"

- **No. of tables: 2**
- **No. of figures: 6**
- **No. of pages: 15**

| Table S1. Biocompatible electrochemical media (catholyte) for *C. necator* growth. | |
| --- | --- |
| **Nutrients** | **Concentration** |
| MgSO_4_ | 0.5 g L^-1^ |
| K_2_SO_4_ | 0.5 g L^-1^ |
| Na_2_HPO_4_ | 1 g L^-1^ |
| Bicarbonate Buffer | 50 mM |
| (NH_4_)Cl | 1 g L^-1^ |
| CaCl_2_ | 0.047 g L^-1^ |
| Trace minerals (1000x) | 1 mL L^-1^ |

| **Table S2.** Trace minerals solution for biocompatible media. | |
| --- | --- |
| **Minerals (1000x stock)** | **Concentration** |
| CuSO_4_·5H_2_O | 0.480 g L^-1^ |
| ZnSO_4_·7H_2_O | 2.400 g L^-1^ |
| MnSO_4_·H_2_O | 2.400 g L^-1^ |
| FeSO_4_·7H_2_O | 15.000 g L^-1^ |

**Text S1. Bioreactor configuration**

Bioreactor experiments were conducted using a bioXplorer® 400P system equipped with 8 parallel stirred-tank reactors (HEL Ltd., UK). Each reactor had a total volume of 500 mL and was operated with a typical working liquid volume of 300 mL. Reactor operation, data acquisition, and process control were performed using WinIso software. Each bioreactor was fitted with two gas inlets connected to gas spargers for CO_2_/N_2_/O_2_. Individual gas streams were regulated using mass flow controllers, and total inlet flow rates were adjusted as required to maintain the desired gas composition. The reactors were equipped with probes and control systems for pH, dissolved oxygen (DO), agitation, temperature, and pressure. A condenser was installed on the exhaust line to minimize evaporative water loss during operation. Liquid samples were withdrawn periodically through a sterile sampling port fitted with a two-way valve. All experiments were conducted at 30 °C. Reactor pH was controlled automatically by addition of 14% (w/w) ammonium hydroxide. Dissolved oxygen was maintained using a cascade control strategy in which agitation increased when DO dropped below the set point of 10%.

**Text S2. Performance metrics and statistical analysis**

Formate selectivity is reported as coulombic efficiency (*CE*), representing the fraction of electrons directed toward formate formation. *CE%* for formate production was calculated based on the theoretical amount of charge ($Q_{{HCOO}^{-}}$), and total amount of charge ($Q_{total}$) required for formate production:

$$\eta_{{HCOO}^{-}}\left( mmol \right)=C_{{HCOO}^{-}}\times V_{cath}\times1000$$

where $\eta_{{HCOO}^{-}}$ are moles of formate produced, $C_{{HCOO}^{-}}$ is the concentration of the formate produced (mol L^-1^), and ($V_{cath}$) is the catholyte volume (L).

$$CE \left( \% \right) or Selectivity to Formate= \frac{Q_{{HCOO}^{-}}}{Q_{total}}=\frac{\eta_{{HCOO}^{-}}\mathcal{\times z\times}F}{\int_{t=0}^{t} Idt}\times100$$

where 𝑧 = 2 is the electron stoichiometry for formate, 𝐹 is Faraday’s constant (96485 C mol^−1^), 𝐼 is the current (A) and 𝑡 is the time (s).

The rate of formate production was calculated based on the change in formate concentration over time and normalized to cathode surface area (A):

$$r_{HCOO-}\left( \frac{mmol}{hr {cm}^{2}} \right)=\frac{\eta_{{HCOO}^{-}}}{A\times dt}$$

Carbon conversion efficiency (CCE) was calculated as the fraction of carbon fed as CO₂ that was recovered as carbon in formate:

$${CCE}_{HCOO-}\left( \% \right)=\frac{\eta_{C-{HCOO}^{-}}}{\eta_{C-{CO}_{2}}}$$

Since formate contains one carbon atom per mole, carbon in formate equals moles of formate produced:

$$\eta_{{HCOO}^{-}}=\eta_{C-{HCOO}^{-}}$$

The moles of CO₂ fed to the reactor were calculated from the inlet volumetric flow rate using the ideal gas law:

$$\eta_{{CO}_{2, in}}=\frac{\left( P\times V_{{CO}_{2}}\times t \right)}{R\times T}$$

where *P* is pressure (Pa), V_CO2_ is the CO_2_ volumetric flow rate (m^3^ S^-1^), t is time (s), R is the ideal gas constant (8.314 J mol^-1^ K^-1^), and T is temperature (K). Thus,

$${CCE}_{{HCOO}^{-}}\left( \% \right)=\frac{\eta_{{HCOO}^{-}}}{\eta_{{CO}_{2}, in}}\times100$$

Energy consumption per mole of formate ($P_{formate}$, kWh mol⁻¹) is typically calculated from the electrical energy supplied to the reactor divided by the number of moles of formate produced.

$$P_{formate}\left( kWh {mol}^{-1} \right)=\frac{\left( V\times I\times t \right)}{1000\times\eta_{{HCOO}^{-}}}$$

Where V is average cell voltage (V), I is the current (A), t is the operation time (h), and $\eta_{{HCOO}^{-}}$is the formate produced (mol).

All experiments were performed in duplicate, and results are reported as mean ± standard deviation. To assess relationships between operating parameters and system performance, correlation analysis was conducted using Spearman’s rank correlation coefficient (ρ), where correlation values range from −1 to +1. Multivariate patterns in the dataset were further explored using principal component analysis (PCA), and results were visualized as biplots generated in R (RStudio, version 1.4.1103).

**Text S3. Energy Efficiency and Power Input.**

The energy efficiency of the CO₂-to-formate conversion process was assessed by determining the energy consumption (*P*_formate_) in kilowatt-hours per mole of formate (kWh mol-FA^-1^) under varying cathodic potentials and fluidized catalysts **(Fig. S1)**. The results demonstrate the critical impact of both potential and catalyst type on the energy efficiency of the system. For the Bi fluidized system, the lowest P_formate_ of 0.12 kWh mol-FA^-1^ was achieved at -2 V, signifying optimal energy efficiency at this potential. At less negative potential of -1.5 V, P_formate_ was slightly increased to 0.13 kWh mol-FA^-1^, while at -1.2 V, P_formate_ increased significantly to 0.18 kWh mol-FA^-1^. Similarly, the Sn fluidized system showed a P_formate_ of 0.12 kWh mol-FA^-1^ at -2 V and -1.5 V. However, at -1.2 V, energy consumption rose sharply to 0.22 kWh mol-FA^-1^. The increase in energy consumption at lower overpotentials aligns with reduced selectivity for formate production as competing reactions, such as HER, become more dominant. The In fluidized system also revealed its best energy performance at -1.5 V, with a P_formate_ of 0.12 kWh mol-FA^-1^, slightly outperforming Bi and Sn under similar conditions. However, at -2 V and -1.2 V, P_formate_ values increased to 0.13 and significantly to 0.24 kWh mol-FA^-1^, respectively, suggesting a less favorable energy profile at these potentials. The flat electrode system without fluidized catalyst exhibited moderate energy efficiency, with P_formate_ values of 0.16, 0.13, and 0.17 kWh mol-FA^-1^ at -2 V, -1.5 V, and -1.2 V, respectively. While reasonable efficiency has been demonstrated at -1.5 V, it was outperformed by the configurations using fluidized catalyst systems across all potentials.

These findings emphasize the importance of operating potential and catalyst selection in optimizing energy efficiency for the CO_2_-to-formate conversion process. The Sn and Bi fluidized system at -2 V, or the In at -1.5V, provided the best combination of low energy consumption and high formate selectivity, making it an ideal candidates for scaling up energy-efficient electrosynthesis systems. From a scalability perspective, achieving high energy efficiency is critical to minimizing operational costs and environmental impacts. The high-power consumptions at -2 V were balanced by significant gains in formate production efficiency, making it a feasible operating point for practical applications.

Future studies should explore the integration of renewable energy sources to offset the high-power input associated with more negative applied potentials. Additionally, life-cycle assessments could provide a more comprehensive understanding of the energy and environmental implications of the CO_2_-to-formate electrochemical process.

**Text S4. Principal Component Analysis of CO₂ Reduction Performance.**

To systematically assess how key operating variables, interact to influence overall system performance, we employed principal component analysis (PCA). This multivariate statistical tool simultaneously considered variables such as applied potential, catalyst material, current density, and Faradaic efficiency to uncover underlying correlations between operating conditions and electrochemical outcomes **(Fig. S2).** Performance parameters, including current density, formate production rate, and *CE*, align strongly along the positive PC1 axis. This clustering means that these metrics are positively correlated, reflecting that higher formate production and *CE* are associated with increased current density. In contrast, HER is negatively correlated with these metrics, as seen in its position along the negative PC1 axis. Energy efficiency aligns positively with PC1 but shows slight variability compared to other formate-related metrics, indicating its distinct but related contribution to overall system performance.

In terms of cathode materials and applied potentials, the electrocatalysts Sn, Bi, and In at -2 V are clustered in the positive PC1 region, strongly aligned with high *CE*, current density, and formate production rates. This confirms that high negative potentials (-2 V) enhance CO_2_ reduction efficiency and selectivity for formate production. Among these, the -2 V with Sn catalyst condition exhibits the highest performance, closely associated with the strongest alignment with the primary performance metrics. The conditions of -1.5 V using Bi and Sn catalysts are positioned between the high- and low-performance regions. These points reflect moderate formate production and *CE*, highlighting the trade-off between applied potential and catalytic activity. However, the condition of -1.2 V and with either In, Sn, Bi or without particles, all cluster in the negative PC1 region, showing higher HER activity and lower energy efficiency. As expected, the less negative potential diminishes the selectivity for CO_2_-to-formate conversion, favoring HER instead. The system without catalytic particles, regardless of the applied potential, exhibit poor performance, as evidenced by their clustering in the negative PC1 region. This finding underscores the necessity of the fluidized electrocatalyst to achieve efficient CO_2_ reduction and selectivity to formate, as systems without particles perform poorly across all conditions.

These findings emphasize the importance of optimizing both the applied potential and catalyst selection to achieve efficient and selective CO_2_-to-formate reduction process. The insights provided by this analysis contribute to the rational design of advanced electrosynthesis cell for sustainable formate production.

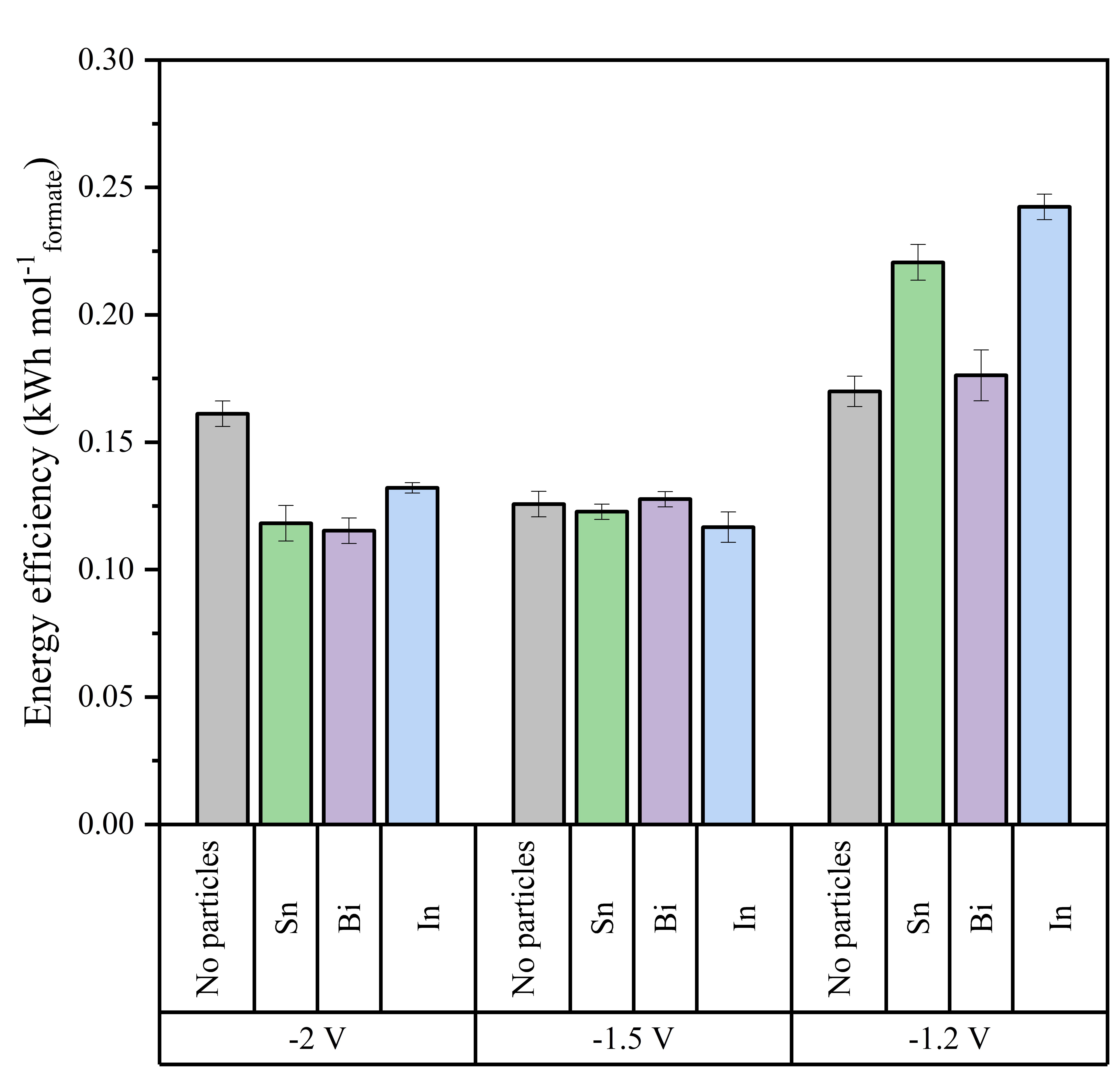

**Figure S1.** Energy efficiency with different electrocatalysts under different applied potentials. Experiments were performed in duplicate, and results are reported as mean ± standard deviation.

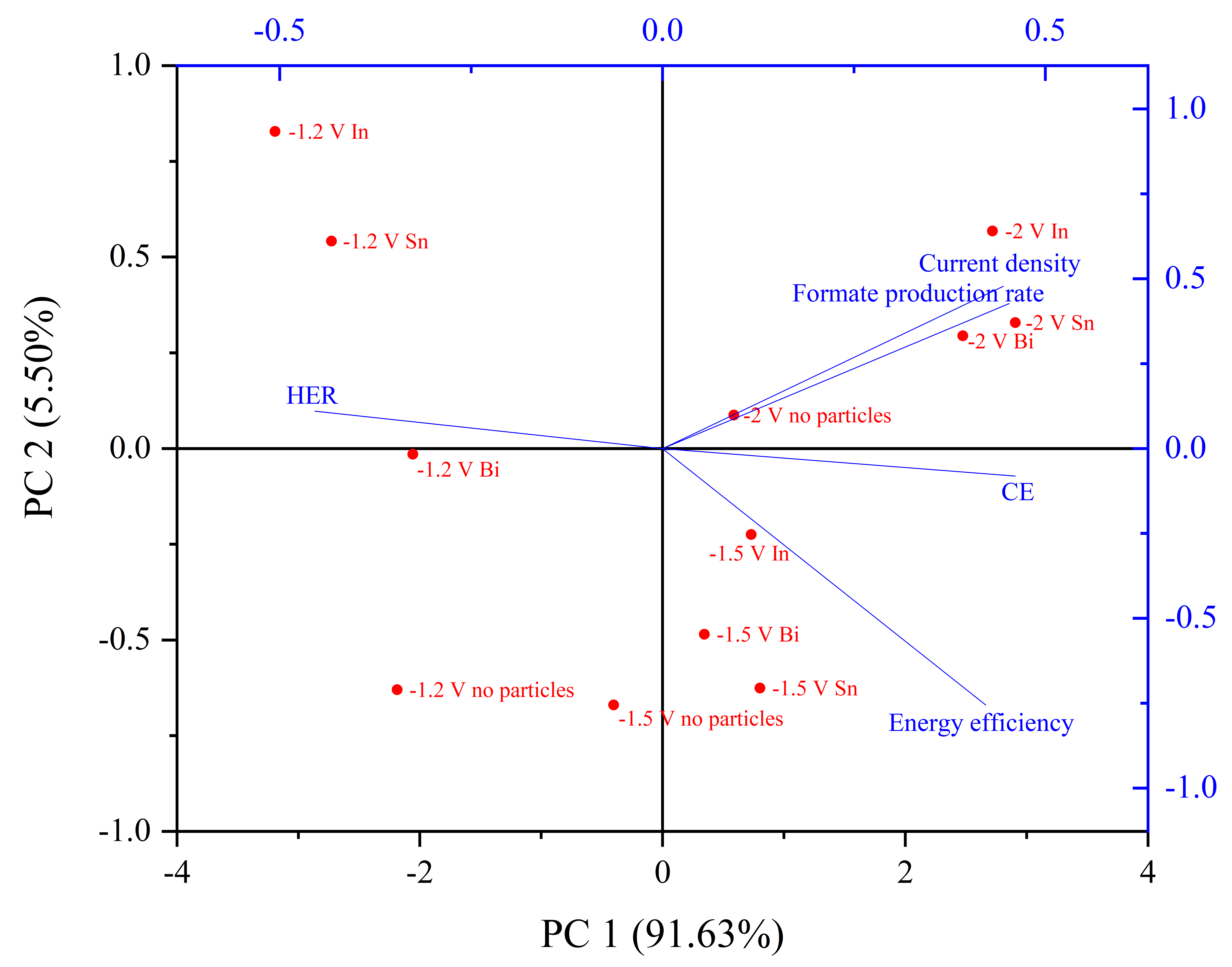

**Figure S2.** Multivariant analysis to explore the impact of different parameters on CO_2_-to-formate efficiency.

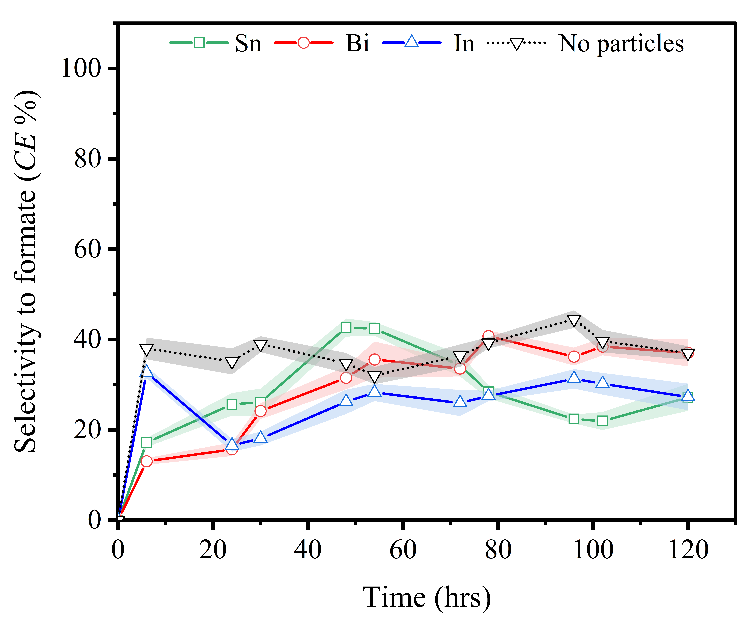

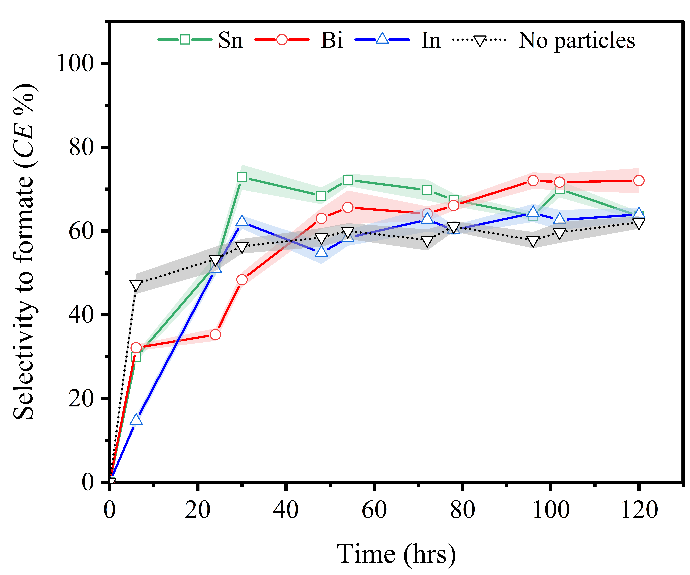

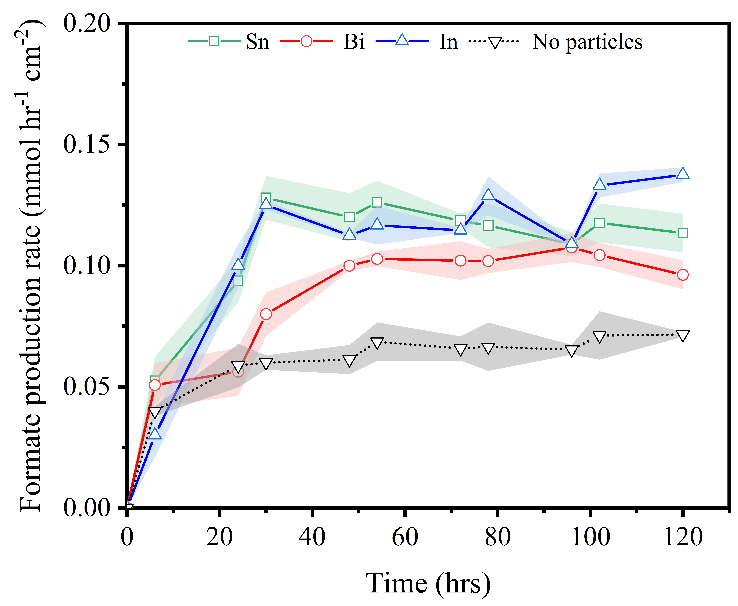

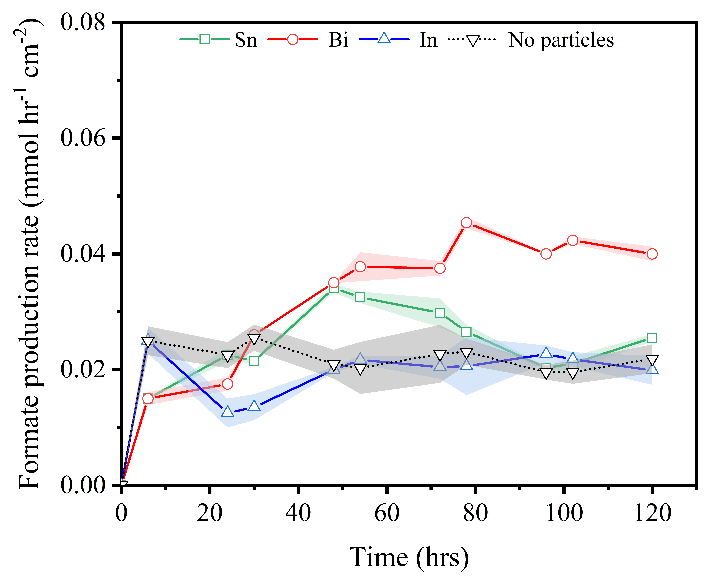

(A)

(B)

(C)

(D)

**Figure S3.** Influence of electrocatalyst materials on CO_2_-to-Formate efficiency. (A-B) at -1.5 V, and (C-D) at -1.2 V vs. Ag/AgCl. Experiments were performed in duplicate, and results are reported as mean ± standard deviation (shown with a shaded area).

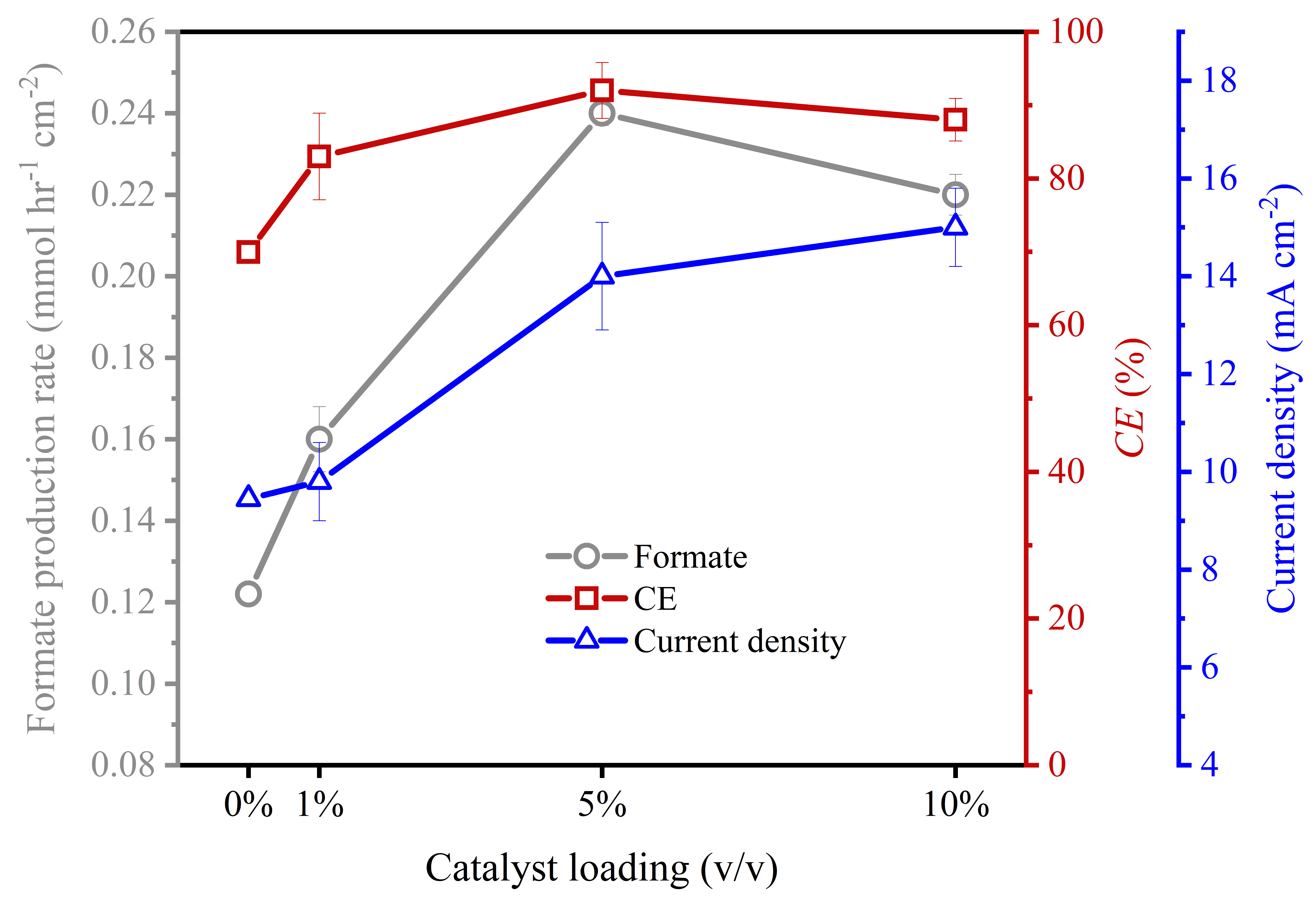

**Figure S4.** Effect of catalyst loading on CO_2_ reduction performance. Sn catalyst at -2 V vs. Ag/AgCl was used. Experiments were performed in duplicate, and results are reported as mean ± standard deviation.

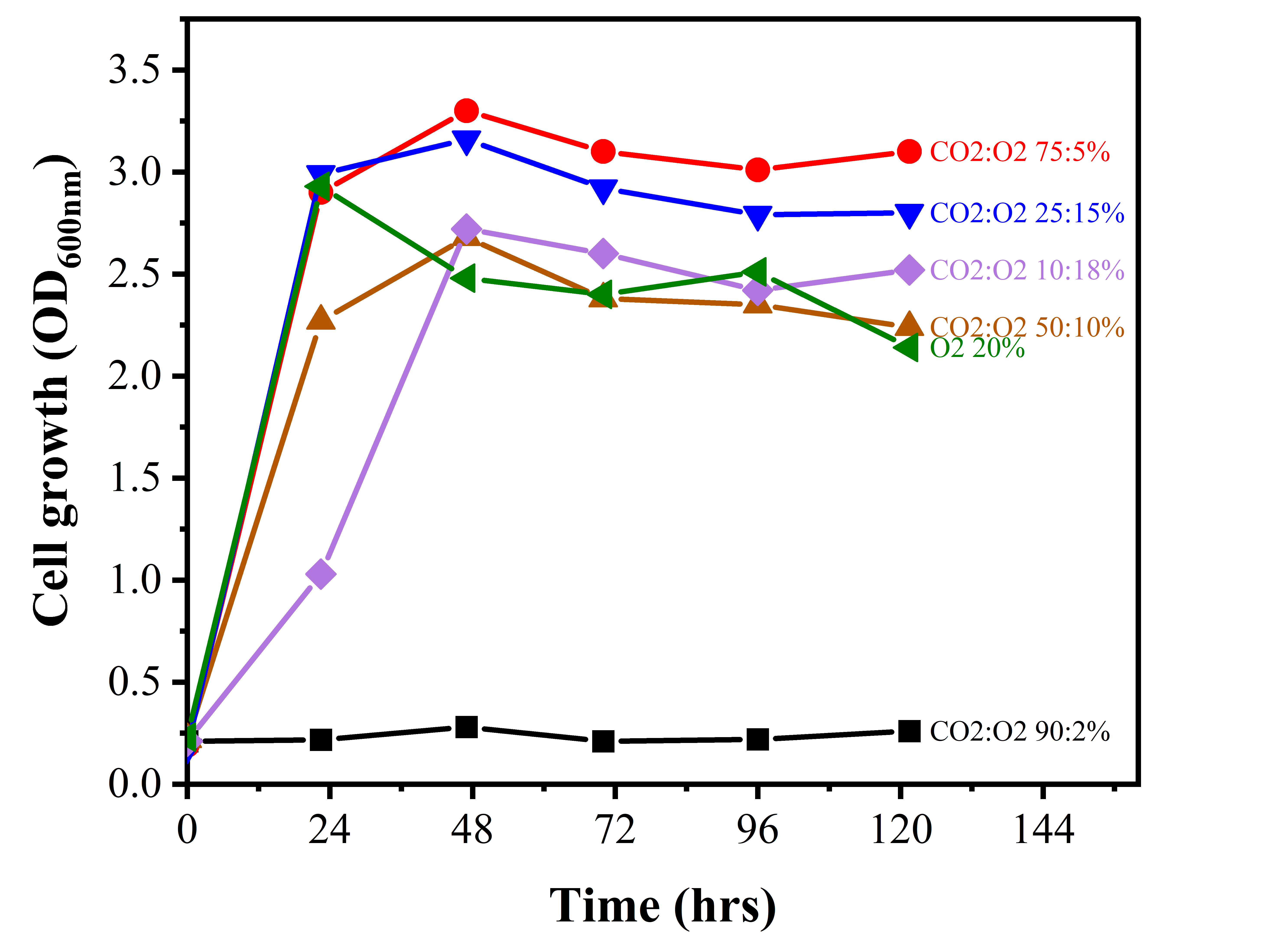

.

**Figure S5.** *C. necator* growth under aerobic conditions with different CO_2_:O_2_ mixtures. This experiment was performed in bioreactors in biocompatible media supplemented with glucose 5g L^-1^ to determine ideal gas mixture for aerobic biohybrid experiments.

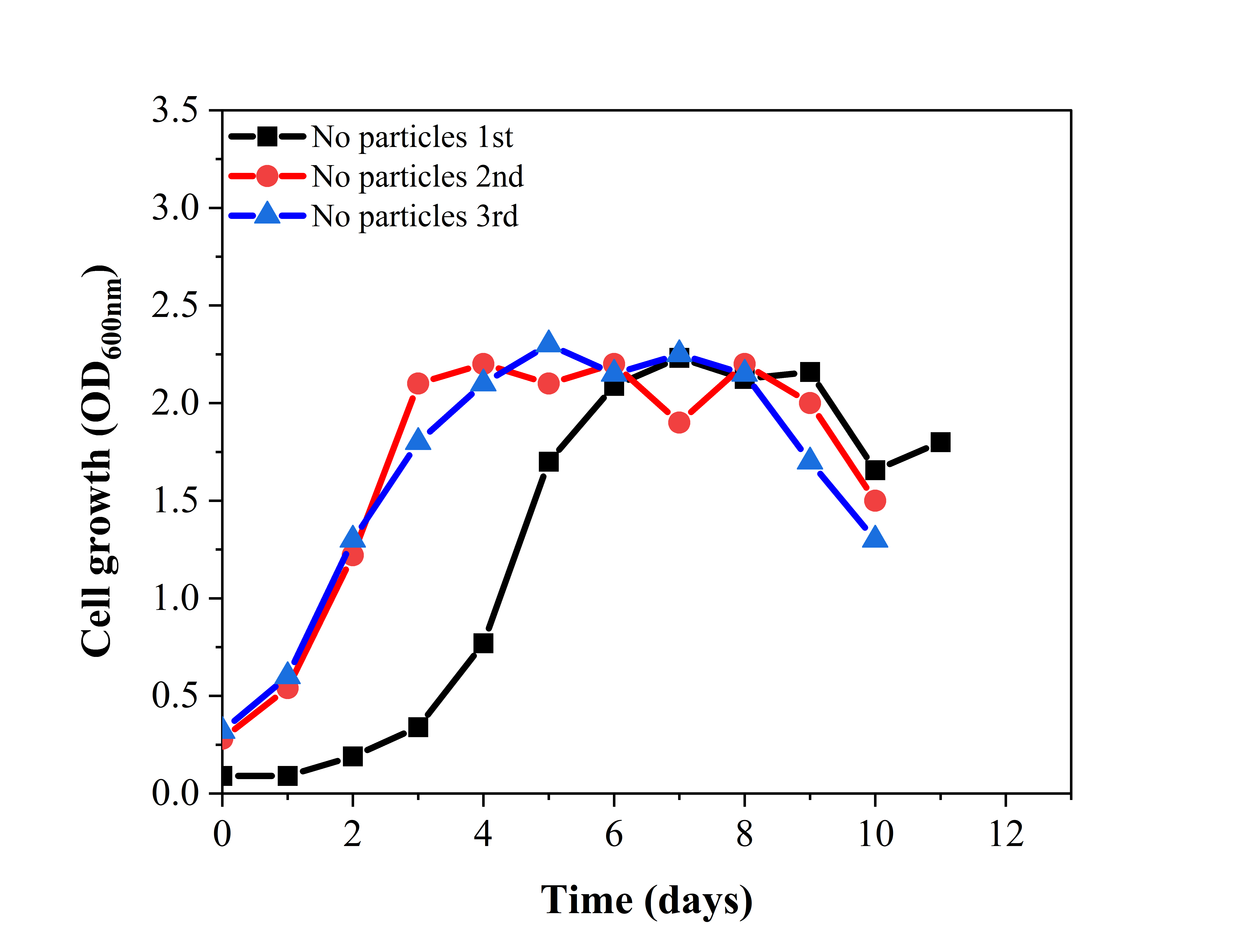

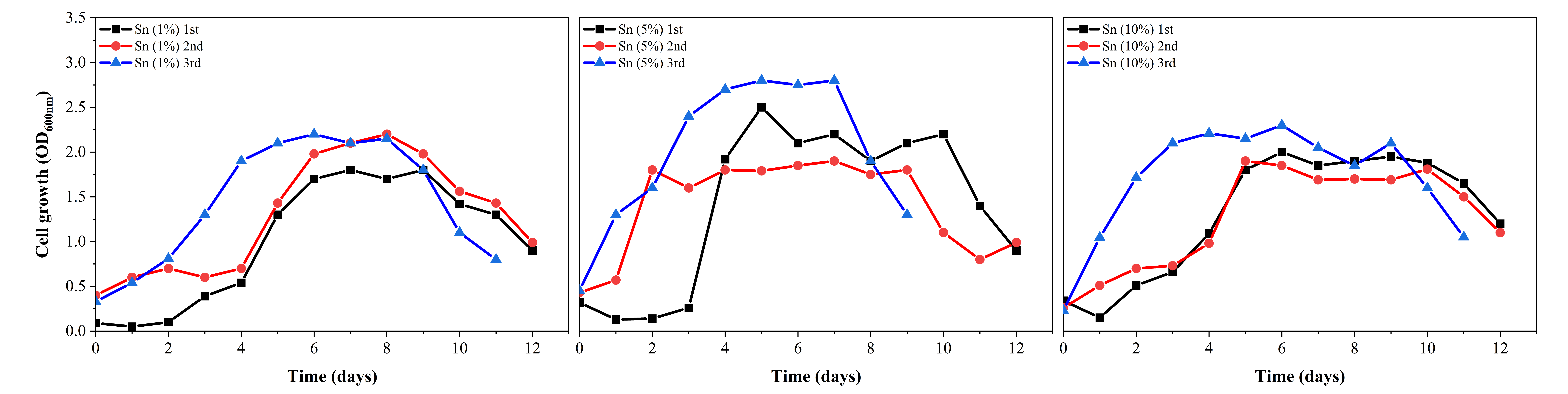

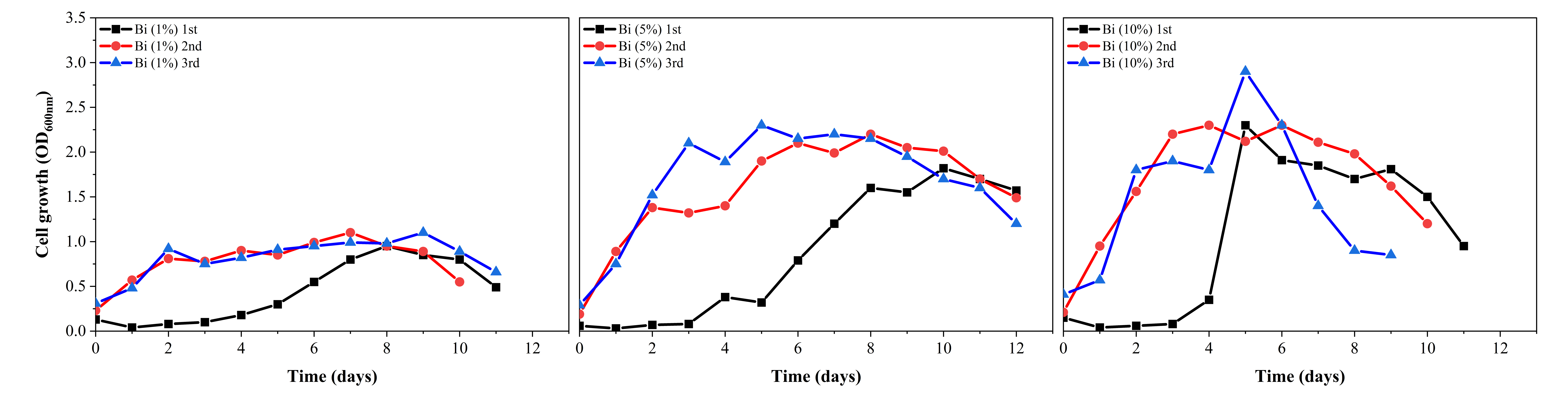

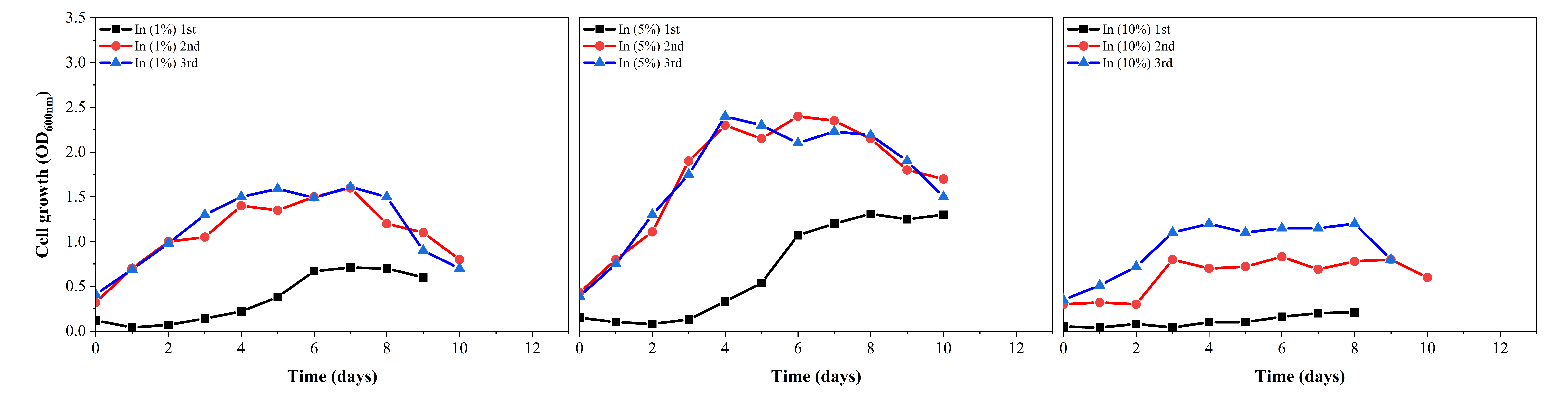

(A)

(B)

(C)

(D)

**Figure S6.** Acclimation of *C. necator* growth in the biocompatible media with electrocatalysts at different loadings in shaken flasks. The curves show microbial growth after several transfers. The inoculum used for the 2^nd^ transfer (2^nd^) was growth medium of the flask of the 1^st^ transfer (1^st^), and the inoculum used for the 3^rd^ transfer (3^rd^) was growth medium of the flask of the 2^nd^ transfer. (A) Sn particles, (B) In particles, (C) Bi particles, and (D) with no particles.
